# Poplar CLE peptides stimulating ectomycorrhizal symbiosis uncovered by genome-wide identification of ectomycorrhizal-responsive small secreted peptides

**DOI:** 10.1101/2025.04.22.649968

**Authors:** Clémence Bonnot, Emmanuelle Morin, Alexandra Henocq, Alexis Chartoire, Emilie Da Silva Machado, Claire Veneault-Fourrey, Annegret Kohler, Francis Martin

## Abstract

Plants small secreted peptides (SSPs) regulate root development, immunity and symbiotic relationships in herbaceous plants. These processes are equally important for establishing functional ectomycorrhizal associations in trees. While fungal SSPs involved in ectomycorrhizal establishment have been identified, the role of plant SSPs remains largely unexplored. Although thousands of SSPs were predicted in plant genomes, their small size and high sequence divergence hinder their accurate automated annotations. To address this issue, we combined *de novo* gene prediction with family-specific motif search to identify 1,053 SSPs from 21 symbiosis-related families in the genomes of two ectomycorrhizal tree species: *Populus trichocarpa* and *Quercus robur*. Nearly half of these SSPs are transcriptionally regulated during ectomycorrhizal symbiosis with various fungal partners. Functional assays of selected *Populus* CLE peptides, a SSP family known to repress arbuscular mycorrhizal and rhizobial symbioses, revealed that five enhanced ectomycorrhizal roots formation. Unlike CLEs involved in the autoregulation of arbuscular mycorrhizal and rhizobial symbioses, these peptides belonged to clades associated with meristem activity. Their activity did not increase lateral root number but inhibited adventitious root growth, suggesting their role in promoting ectomycorrhizal root organogenesis. These findings demonstrate that CLE peptides promote, rather than repress, ectomycorrhizal symbiosis. This functional divergence from their roles in other symbioses suggests that poplar co-opted a distinct set of SSPs to support ectomycorrhizal development. Our results expand the understanding of host tree contributions to ectomycorrhizal development and identify a set of candidate SSPs for future functional studies, thereby highlighting a new layer of regulation in tree-fungal mutualism.

## Introduction

In boreal and temperate forests, most trees form symbiotic associations with ectomycorrhizal (ECM) fungi to ensure nutrition. Providing mineral nutrients and water in return for organic carbon released by their tree host, ECM fungi contribute to tree growth and health, and are pivotal for carbon sequestration and nutrient cycling in these ecosystems (van Der Heijden et al., 2015; Martin et al., 2016; Steidinger et al., 2019). Over the last 20 years, the increase in available genomes of ECM tree species such as poplar (*Populus trichocarpa;* (Tuskan et al., 2006)), eucalypt (*Eucalyptus grandis;* (Myburg et al., 2014)) or oak (*Quercus robur;* (Plomion et al., 2018)), and fungal species (1000 fungal genomes; (Grigoriev et al., 2014)), coupled with the development of *in vitro* ECM models such as *Populus × canescens* associated with *Laccaria bicolor,* and *E. grandis* associated with *Pisolithus microcarpus,* engaged in the dissection of the molecular mechanisms leading to ECM associations (Marqués-Gálvez et al., 2022; Plett et al., 2024a). *In vitro* experiments associated with transcriptomics identified several fungal metabolites (Ditengou et al., 2003; Ditengou et al., 2015; Felten et al., 2009; Vayssières et al., 2015; Plett et al., 2024b), peptide effectors (Plett et al., 2011; Plett et al., 2014a; Plett et al., 2020b; Kang et al., 2020; Marqués-Gálvez et al., 2024), and one miRNA (Wong-Bajracharya et al., 2022), whose interferences with various plant metabolic pathways are necessary for the establishment and maintenance of ECM symbiosis. However, no common ECM gene toolkits were identified in fungi or trees (Kohler et al., 2015; Miyauchi et al., 2020; Marqués-Gálvez et al., 2022; Marqués-Gálvez et al., 2025). Cope et al. demonstrated the role of elements of the plant common symbiosis pathway in the establishment of the ECM association between poplar and *L. bicolor* (Cope et al., 2019). Controlling the establishment of arbuscular mycorrhizal (AM) symbiosis, the common symbiosis pathway is highly conserved in plant species that form these associations, including poplars (Oldroyd, 2013). However, it is not present in species that are unable to establish AM symbiosis, such as most ECM tree species (Garcia et al., 2015; Radhakrishnan et al., 2020). Hence, this pathway is unlikely to be a common component in the establishment of ECM interactions. A recurring feature of ECM accommodation is the attenuation of host tree defence mechanisms. Genes associated with plant defence are downregulated at the transcriptional level during ECM symbiosis, but are induced during non-compatible ECM interactions, or in trees growing under conditions of high-nutrient availability in which ECM associations are repressed (Plett et al., 2014b; Hortal et al., 2017; Basso et al., 2020; Plett et al., 2020a; Marqués-Gálvez et al., 2025). Moreover, during the ECM associations of *P. x canescens* with *L. bicolor* and of *E. grandis* with *P. microcarpus*, plant defence-associated hormonal pathways (respectively the jasmonic acid signalling and the ethylene biosynthesis pathways) seem targeted by fungal peptide effectors (Plett et al., 2014a; Plett et al., 2020b; Daguerre et al., 2020; Marqués-Gálvez et al., 2024). Finally, QTL analyses identified in polar two markers of ECM receptivity associated with plant defences: an Ethylene Responsive Binding Protein Transcription factor known to repress ethylene-associated pathogenesis genes (Labbé et al., 2011), and a G-type lectin Receptor-Like Kinase which permits ECM colonisation of the root apoplast, probably through the repression of plant immunity (Labbé et al., 2019). Their modes of action are unknown.

Small secreted peptides (SSPs) (<250 amino acids) are produced from short open reading frames or through proteolytic cleavage of longer pre-proteins and secreted into the apoplast (Matsubayashi, 2014; Tabata and Sawa, 2014; Tavormina et al., 2015; Hu et al., 2021; Zhang et al., 2025). They belong to various phylogenetic families. Extensive genome and proteome searches uncovered over 4,000 putative SSP families in plants, of which only 69 were functionally characterised (Ghorbani et al., 2015; Tavormina et al., 2015; de Bang et al., 2017; Segonzac and Monaghan, 2019). Ranging from antimicrobials to signalling peptides, they are involved in a wide variety of developmental processes and adaptive plant responses to the environment (Matsubayashi, 2014; Tavormina et al., 2015; Hu et al., 2021; Zhang et al., 2025). Several SSP families encoding signalling peptides involved in intercellular communication (signalling-SSPs) mediate local and long-distance signals that regulate symbiotic associations with Rhizobiaceae bacteria and AM fungi (de Bang et al., 2017; Roy and Müller, 2022). Among these, the C-TERMINALLY-ENCODED PEPTIDE (CEP) and CLAVATA3/EMBRYO SURROUNDING REGION-related (CLE) families are those whose roles and modes of action in symbiosis are best characterised. During nitrogen and phosphorus deficiencies, the secretion of CEP peptides in the xylem sap transduces root-to-shoot starvation signals, leading to the systemic promotion of nodulation (Imin et al., 2013; Mohd-Radzman et al., 2016; Gautrat et al., 2020; Laffont et al., 2020, 202; Zhu et al., 2020a) and AM associations (Pedinotti et al., 2024). In contrast, the secretion of CLE peptides conveys both local and systemic satiety signals inhibiting nodulation and AM associations in response to nitrate and inorganic phosphate, but also to prior associations with AM and Rhizobiaceae bacteria (Okamoto et al., 2008; Mortier et al., 2010; Reid et al., 2011; Nishida et al., 2016; Müller et al., 2019; Karlo et al., 2020; Mens et al., 2021; Wang et al., 2021a; Wulf et al., 2024). At the site of nodule initiation, the transcriptional regulation of peptides of the GLV/RGF/CLEL (GOLVEN/ROOT GROWTH FACTOR/CLE-LIKE) family represses local nodule organogenesis in response to nitrate and rhizobial symbiosis (Li et al., 2020; Li et al., 2023; Roy et al., 2024). Members of the PSK (PHYTOSULFOKINE), DVL/RTFL (DEVIL/ROTUNDIFOLIA), and RALF (RAPID ALKALINIZATION FACTOR) families regulate early-stages of rhizobial infection and late nodule development (Combier et al., 2008; Wang et al., 2015; Di et al., 2022; Yu et al., 2022). The transcriptional regulation of members of 22 other SSP families during rhizobial or AM symbiosis suggests that more SSPs families are involved in these symbioses (Massoumou et al., 2007; van Wyk et al., 2014; de Bang et al., 2017). In the context of ECM symbiosis, 417 putative SSPs were found transcriptionally upregulated in poplar during its ECM association with *L. bicolor* (Plett et al., 2017). One of them was shown to penetrate fungal nuclei where it interferes with a fungal transcription factor. Its over-expression was associated with a 10% increase of ECM root formation demonstrating its participation to the ECM process (Liu et al., 2024). Despite these promising findings, the broader role of plant SSPs in ECM symbiosis remains largely unexplored.

To investigate whether more SSPs in trees mediate the formation or regulation of ECM associations, we focused on 21 SSP families related to symbiosis in herbaceous plants (Sup. Table S1) and encoded in tree genomes (Philippe et al., 2009; Jiménez-López et al., 2011; Onrubia et al., 2014; Ghorbani et al., 2015; Guo et al., 2015; Cao et al., 2015; Furumizu et al., 2021; Wang et al., 2021b; Wang et al., 2023). All were selected based on their known participation in the regulation of rhizobial or AM symbiosis (CLE, CEP, DVL, GLV, PSK, RALF, nsLTP, PCY and PHYCYS) (van Wyk et al., 2014; Sun et al., 2019; Roy and Müller, 2022; Chen et al., 2023), or on their transcriptional regulation during these symbiotic associations (CAPE, ECL, EPFL, GASA, GRI, IDL, KTI, PIPL, POE, PSY, TAX and TPD) (Massoumou et al., 2007; de Bang et al., 2017). Most, such as the CLE and CEP families, encode signalling-SSPs that transduce cell-cell signals through the binding of cell-surface receptors (Tavormina et al., 2015; Olsson et al., 2019; Kim et al., 2021; Keerthana et al., 2025), while others encode peptides associated with antimicrobial functions without knowledge of their modes of action (CAPE, GASA and nsLTP) (Breen et al., 2017; Fleury et al., 2019; Gao et al., 2022; Han et al., 2023; Nahirñak et al., 2024) or protease inhibitors (KTI and PHYCYS) (Martínez et al., 2012; Rustgi et al., 2018; Balbinott and Margis, 2022). We established their gene repertoires in two ECM tree species (*P. trichocarpa* and *Q. robur*) and analysed their expression in response to ECM associations with diverse fungal species. We identified 1053 SSPs from the two ECM tree species, of which 40% were differentially expressed during ECM symbiosis. Using synthetic peptides, we showed that five ECM-induced CLE peptides from poplar stimulated ECM association with *L. bicolor*. Our results demonstrate for the first time that a functional role for CLE peptides in promoting ECM symbiosis. This contrasts with their well-established role as negative regulators of AM and rhizobial symbioses (Nakagami et al., 2024). This study also provides a list of SSPs transcriptionally regulated by ECM symbiosis, which is of significant interest for further investigation of the role of SSPs in ECM symbiosis.

## Results

### Identification of 21 AM and rhizobial symbiosis-related SSP families in *P. trichocarpa* and *Q. robur*

Discriminated against by classical gene prediction methods, SSP-encoding genes are underrepresented in genomic annotations (Lease and Walker, 2006; Hanada et al., 2007; Silverstein et al., 2007; Pan et al., 2012; Ghorbani et al., 2015; de Bang et al., 2017). Therefore, to identify the members of the 21 selected SSP families in the genomes of *P. trichocarpa* and *Q. robur,* we built an identification pipeline combining motif similarity searches using HMM and MAST on predicted proteomes, and SSP-targeted *de novo* gene predictions on genomes by both global (blast) and motif (HMM) similarity searches (Fig. 1 a). We tested its effectiveness on *Arabidopsis thaliana* and *Medicago truncatula*, two species whose genomes have been extensively searched for members of these families (among many others: Wen et al., 2004; Silverstein et al., 2007; Martinez and Diaz, 2008; Zhou et al., 2013; Ghorbani et al., 2015; Cao et al., 2015; Tost et al., 2021).

**Fig. 1.**
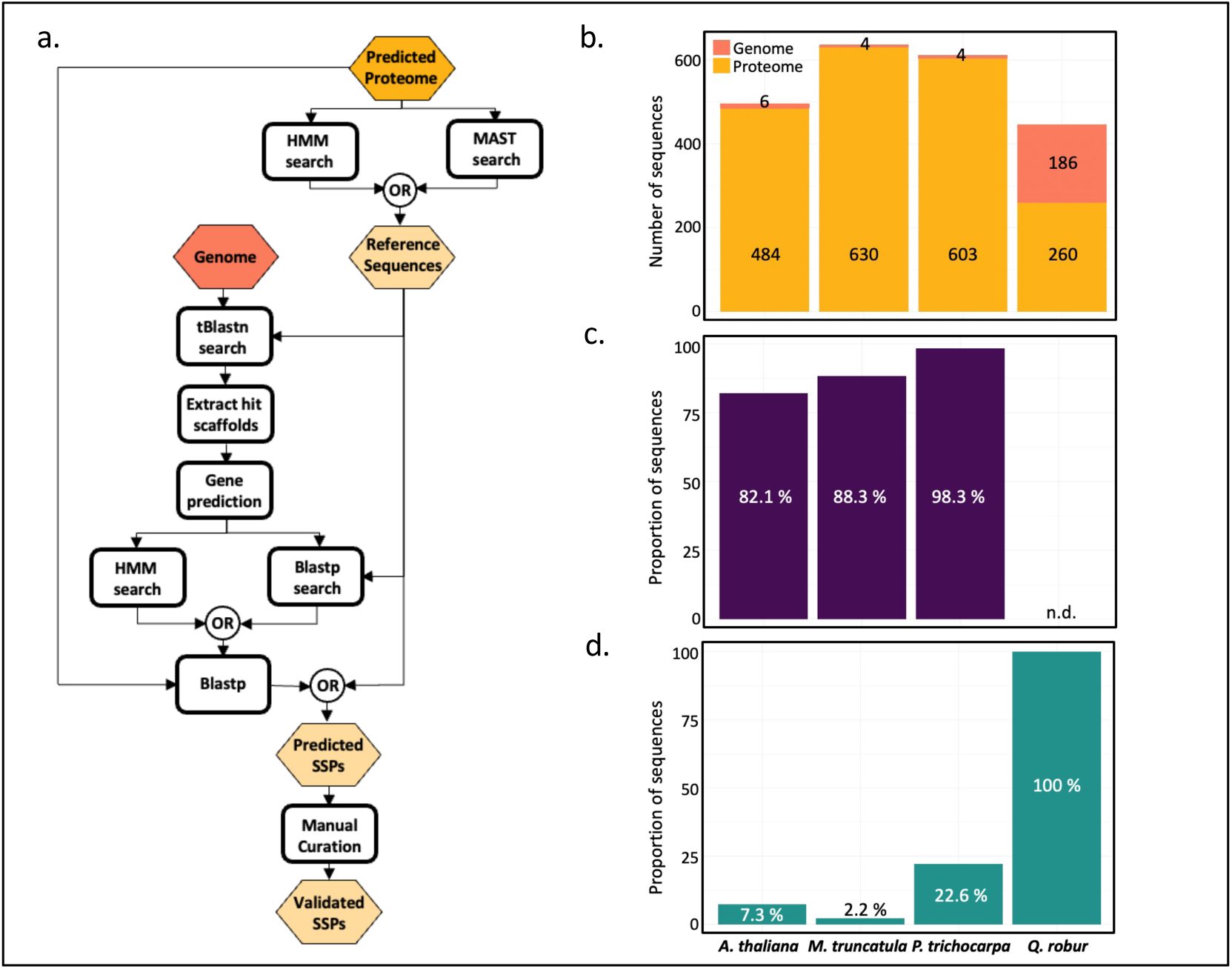
Identification of the gene members of 21 symbiosis-related SSP families in the genome of *A. thaliana*, *M. truncatula*, *P. trichocarpa* and *Q. robur.* a. Identification pipeline (see Materials and Methods for details). b. Total number of SSP sequences identified from the predicted proteome and from the genome of *A. thaliana*, *M. truncatula*, *P. tricocharpa* and *Q. robur*. c. Percentage of the published SSPs recovered from *A. thaliana, M. truncatula, P. tricocharpa* and *Q. robur* by the Identification pipeline. d. Percentage of novel SSPs identified in *A. thaliana*, *M. truncatula*, *P. tricocharpa* and *Q. robur*.

The pipeline identified 2,177 SSPs across the four species (Fig. 1 b, Sup. Table S1, Sup. Table S2). 9.2 % of these (six in *A. thaliana*, four in *M. truncatula*, four in *P. trichocarpa,* and 186 in *Q. robur)* were identified through *de novo* targeted gene prediction (Fig. 1 b, Sup. Fig S1 a, Sup. Table S2). We obtained high recovery rates for the published SSPs. 82.1 % of *A. thaliana’s* published members from 21 families, 88.3 % of *M. truncatula*’s and 98,3 % of *P. trichocarpa’s* were recovered (Fig. 1 c, Sup. Fig S1 b, Sup Table S2). 29 % of the SSPs identified by the pipeline (7.3 % in *A. thaliana*, 2.2 % in *M. truncatula*, 22.6 % in *P. trichocarpa,* and 100 % in *Q. robur*) were novel (Fig. 1 d, Sup. Fig S1 c, Sup. Table S2).

SSP families can be classified into five categories according to their origin and primary sequence characteristics: post-translationally modified (PTM), cysteine-rich (C-rich), functional precursor (FP), short open reading frame (sORF), and those that do not enter any of the previous categories (Other) (Sup. Table 1) (Tavormina et al., 2015). To verify that the identification pipeline was not biased toward any SSP family or category, we used the Spearman rank-order correlation coefficient to test whether the pipeline produced comparable recovery rates of published SSP (Sup. Fig. S2). We found a significant positive correlation for recovery, demonstrating that the pipeline worked similarly for all SSP families and categories. Overall, the large number of new SSPs identified and, the high and consistent recovery rates for all SSP families, suggest that the pipeline is effective and likely identified the 21 SSP families in their entirety for the four plant species.

### The newly identified SSPs share the sequences characteristics of their SSP families

To confirm that the new peptides identified by the pipeline were assigned to the correct families, we verified that they displayed the sequence characteristics of their SSP families. Pairwise similarity clustering of all the SSPs identified by the pipeline showed that novel and published members of the same family clustered together, demonstrating that they shared the strongest full-sequence homologies (Fig. 2 and Sup. Fig. S3). Alignments confirmed the affiliation of nine sequences (three CLEs, three PIPLs, two GLVs, and one PSY) positioned at the outskirts of their families (Sup. Data S1). Maximum likelihood phylogenetic analyses of each of the 21 families produced strong statistical support for all new SSPs sequences, suggesting adequate affiliation (Sup. Fig. S4). PTM and FP SSP families are characterised by the presence of a unique or repeated conserved domain corresponding to mature peptides (Tavormina et al., 2015; Yamaguchi and Kawasaki, 2021; Xie et al., 2022). We used the motif finder software MEME (Bailey and Elkan, 1994) to detect the presence and number of these domains in the peptides of the PTM and FP families (Fig. 3 a and b). We found the typical conserved domains of these families in 98.6% of the sequences. In all families, the occurrence per peptide of the family domain followed the same distribution in all four species (Fig. 3 b). Alignments confirmed the affiliation of eight sequences without family domain (Sup. Data S1). Cysteine-rich families have no conserved domain but a conserved cysteine pattern (Silverstein et al., 2007; Li et al., 2014; Onrubia et al., 2014; Tavormina et al., 2015; Luo et al., 2018). Alignments confirmed the conservation of these patterns in all novel sequences (Sup. Fig. S5, Sup. Data S1).

**Fig. 2.**
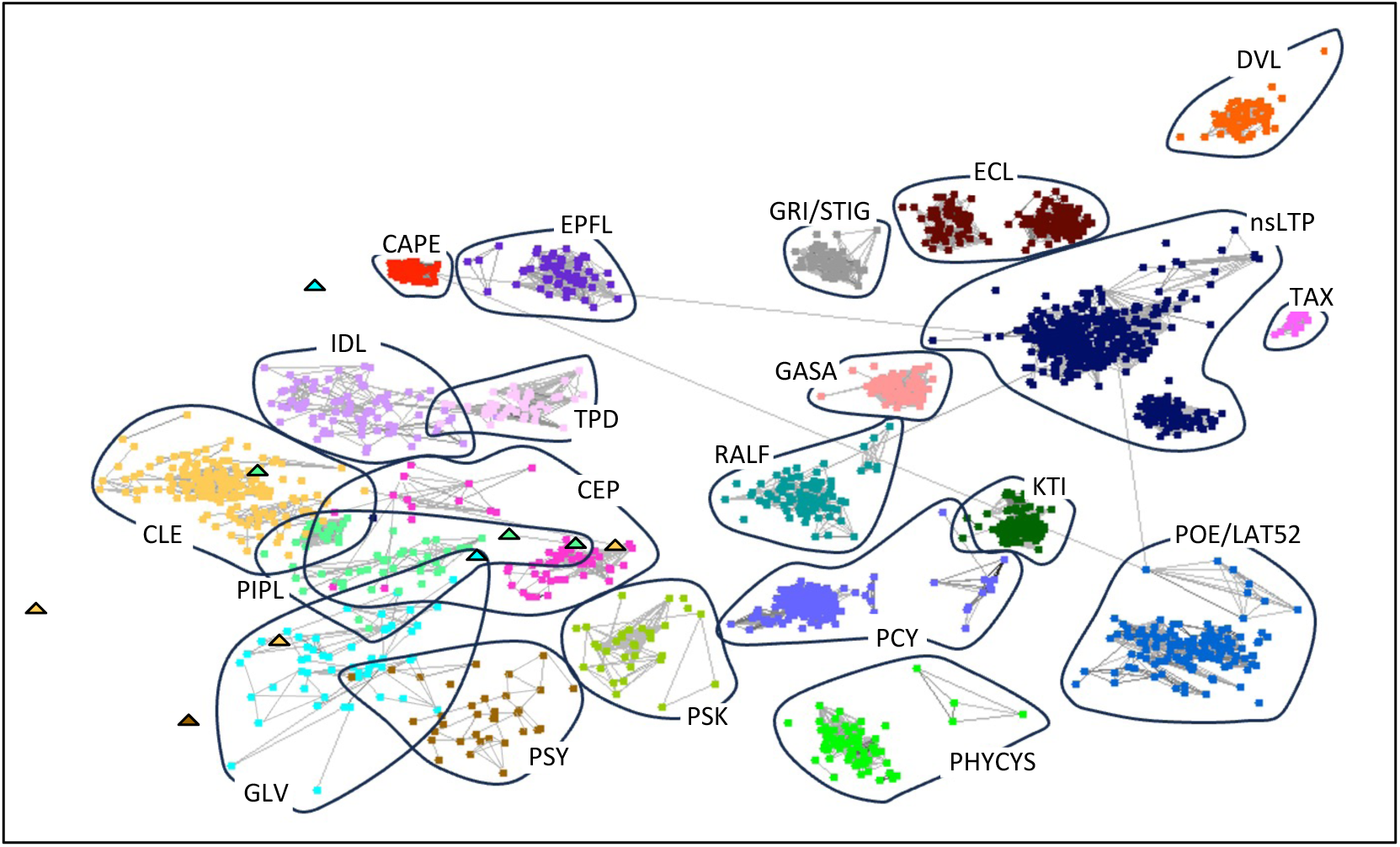
The 2,177 SSP sequences identified in *A. thaliana*, *M. truncatula*, *P. trichocarpa* and *Q. robur* cluster according to their SSP families. Pairwise similarity two-dimensional clustering diagram produced by CLANS (Frickey and Lupas, 2004) of the full amino acid sequences encoded by all identified SSPs. Each amino acid sequence is represented by a square or a triangle. Colours indicate the SSP family of affiliation of each sequence. Sequences are connected by a grey line when the *P-*value of their reciprocal BLASTP hits is P ≤ 10^-10^. The attractive forces between sequences used for clustering are proportional to the negative logarithm of the peptides’ reciprocal BLASTP *P-values*. Triangles mark SSP sequences clustering at the outskirts of their families (the CLEs Medtr1g106920, Medtr7g093050, and Medtr7g084110; the PIPLs AT3G06090, AT3G13370, and prot24979; the GLVs AT5G51451 and Potri10G147100; and PSY AT3G49270).

**Fig. 3.**
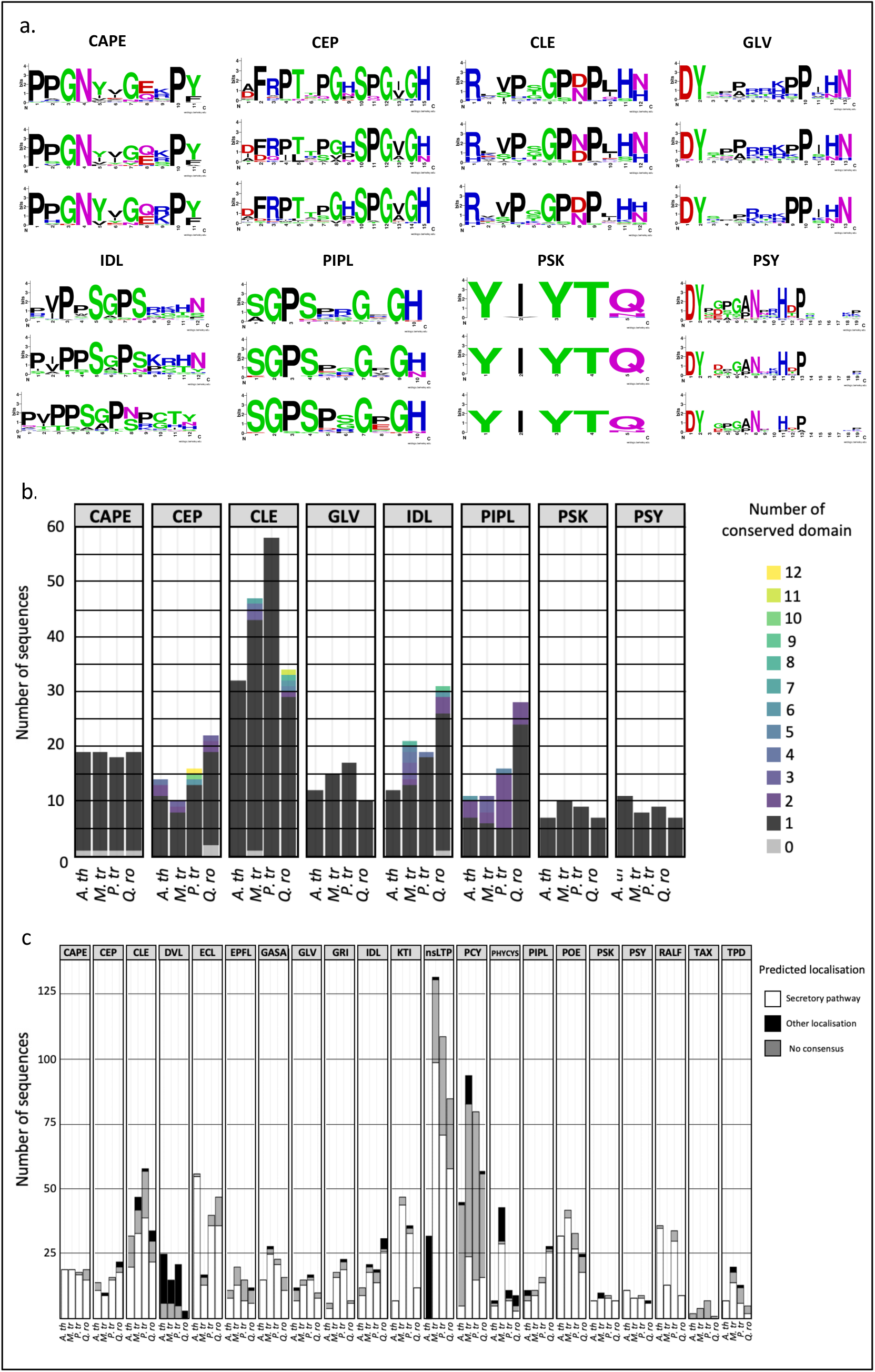
The SSPs identified in *P. trichocarpa* and *Q. robur* showed the typical conserved domains and pattern of predicted location, of their respective families. a. WebLOGO (Crooks et al., 2004) representations of the conserved domain corresponding to the mature SSP in processed SSPs derived from functional precursor (CAPE) or post-translationally modified (CEP, CLE, GLV, IDL, PIPL, PSK and PSY). Domain as found in the herbaceous *A. thaliana* and *M. truncatula* (upper panel) and in the ectomycorrhizal trees *P. trichocarpa* (middle panel) and *Q. robur* (bottom panel). b. Number of conserved domains corresponding to the mature SSPs found per pre-protein in SSPs derived from functional precursors (CAPE) or post-translationally modified (GLV, IDL, PIPL, PSK and PSY). c. Localisation predictions for all members of the 21 SSP families of interest. For each family, the number of sequences predicted by TargetP (Armenteros et al., 2019), SignalP (Teufel et al., 2022), and DeepLoc (Thumuluri et al., 2022) to be sent to the secretory pathway (white), not sent to the secretory pathway (black), or predicted to the secretory pathway by one or two, but not all three prediction softwares (grey, no consensus) are given.

To verify whether the peptides identified by the pipeline were predicted as secreted, we used three subcellular localisation prediction software packages: TargetP (Armenteros et al., 2019), SignalP (Teufel et al., 2022), and DeepLoc (Thumuluri et al., 2022) (Fig. 3 c and Sup. Table S3). 94.4 % of the published peptides and 92.3 % of the novel peptides were predicted to enter the secretory pathway by at least one prediction software. The predicted outcomes were similar for all four species in all families. Among the 21 SSP families, only the DVL family had a majority of sequences not predicted to be secreted. Classified within the SSP families because of their sORF origin, none of the DVL peptides is known to be secreted and act outside the cell (Ikeuchi et al., 2011). Our results showed that the newly identified peptides shared strong global sequence homology with, and the sequence characteristics of, their families, validating their affiliation.

### SSP-encoding genes are transcriptionally regulated by ECM associations in poplar and oak

To assess the involvement of members of the 21 SSP families in ECM symbiosis, we verified the expression of their encoding genes in response to several ECM associations of poplar (*Populus × canescens* with *Laccaria bicolor*, *Cenoccocum geophylum or Pisolithus microcarpus, and Populus tremula × tremuloides with Amanita muscaria)* and oak (*Quercus robur* with *Tuber aestivum*, *Tuber magnatum,* or *Tuber melanosporum)*. We found that 48.6% of poplar and 28.5% of oak genes were regulated by at least one ECM interaction (FDR ≤ 0.05, fold-change ≥ 2) (Fig. 4. a, b, c and d). Most of the regulated SSPs (43 % in poplar and 83.4 % in oak) were repressed by ECM associations (Clusters VIII, IX, X and XI). 30.2 % of poplar- and 14.2 % of oak-regulated SSP genes were induced by ECM associations (Clusters I, II, III, and IV). 26.8 % of poplar and 2.4 % of oak-regulated SSPs genes were oppositely regulated by different ECM interactions (Clusters V, VI, and VII) (Fig. 4. a, b, c, d; Sup. Table S4 and S5).

**Fig. 4.**
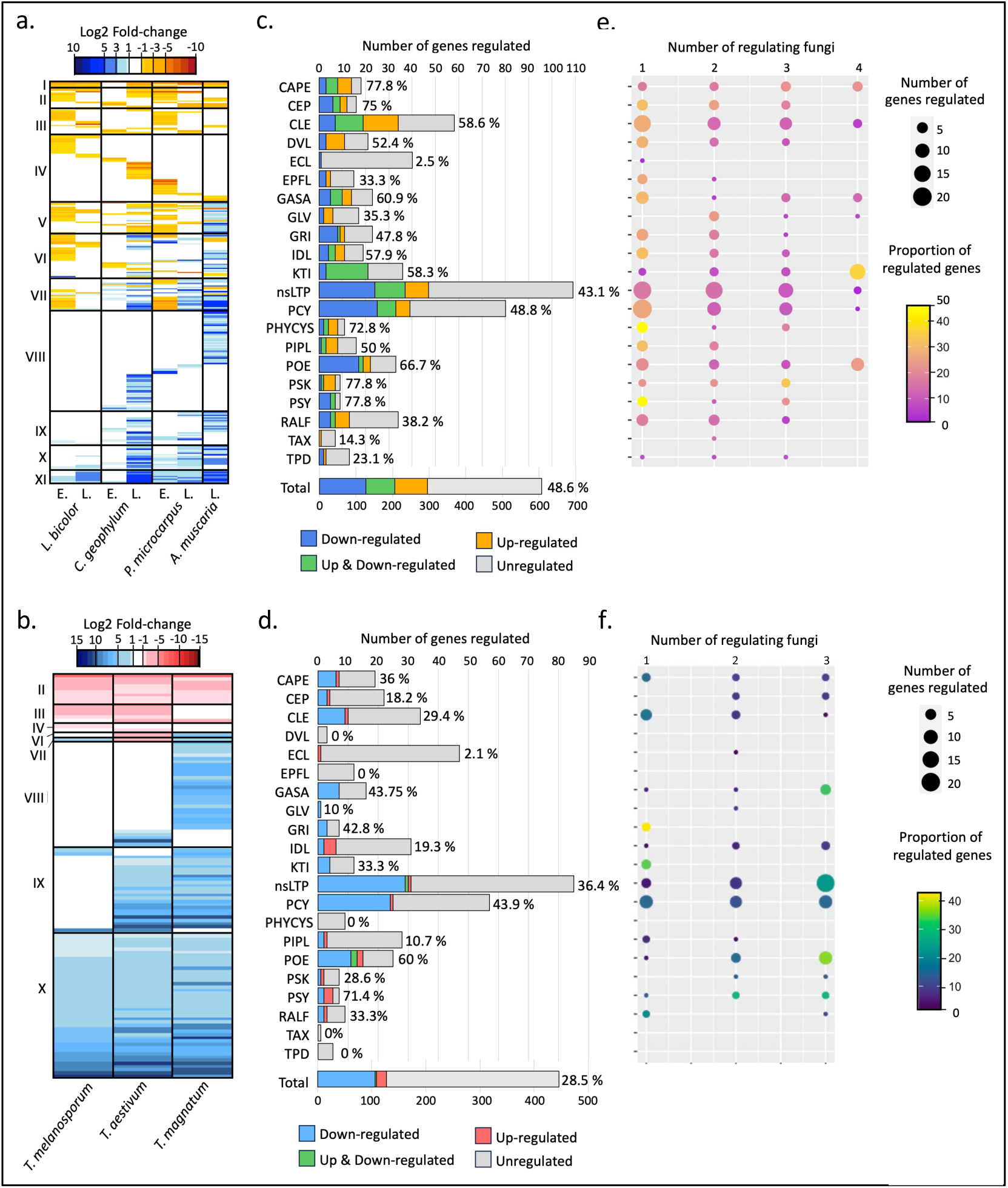
Gene members of all 21 SSP families were regulated in response to ectomycorrhizal symbiosis in poplar and oak. a. and b. Heatmaps representing the clustering of SSP-encoding genes of poplar (*P. tremula **×** alba* for the interaction with *A. muscaria* and *P. **×** canescens* for all others) (a) and oak (*Q. robur*) (b) based on their expression patterns in response to ectomycorrhizal associations with different fungi (log2 fold-change of the ratio of their expression in ectomycorrhiza compared to uncolonized roots). Conditions in which the regulation of a gene was not statistically significant (false discovery rate (FDR) > 0.05) are shown as not regulated (white cells). Genes for which no statistically significant regulation was found for any of the tested conditions are not represented. Roman numbers indicate different gene clusters based on the number of ectomycorrhizal associations that induce or repress the transcription of the SSP-encoding genes. For ease of interpretation, clade names were kept between poplar and oak. Clusters I, II, III, and IV contained SSP-encoding genes induced by four, three, two, or only one ectomycorrhizal interaction, respectively. Clusters V, VI, and VII contained genes oppositely regulated by different ectomycorrhizal interactions: the genes upregulated by more ectomycorrhizal associations in cluster V, the genes up and downregulated by an equal number of ectomycorrhizal associations in cluster VI, and the genes downregulated by more ectomycorrhizal associations in cluster VII. Clusters VIII, IX, X, and XI contained SSP-encoding genes that were repressed by one, two, three, or four ectomycorrhizal associations. a. E. and L. are respectively an early and a late time point of sample collection after contact between the ectomycorrhizal fungi and the host tree (see Material and Method for details). c. and d. Number and proportion of genes of each of the 21 SSP families of interest statistically significantly regulated (upregulated, downregulated or both) by ectomycorrhizal associations in poplar (*P. tremula **×** alba* for the interaction with A. muscaria and *P. **×** canescens* for all others) (c.) and in *Q. robur* (d.). e. and f. Number and proportion of genes of each of the 21 SSP families of interest statistically significantly regulated by one, several, or all ectomycorrhizal fungi tested in association with poplar (*P. tremula **×** alba* for the interaction with A. muscaria and *P. **×** canescens* for all others) (e.) and *Q. robur* (f.).

All 21 SSP families in poplar and 16 in oak possessed at least one member regulated by one of the ECM interactions tested (Fig. 4 c and d). The SSP families with the highest proportion of regulated genes varied between poplar and oak. However, using the Spearman rank-order correlation coefficient, we found that the number of genes regulated by ECM symbiosis was positively correlated with the number of genes in each SSP family in poplar (rs = 0.72, p <0.01) and oak (rs = 0.70, p <0.01) (Sup. Fig. S6), demonstrating that ECM symbiosis regulates similar proportions of genes in all SSP families, with no differences between families or their modes of action (Sup. Table S1). 13 SSP families possessed members that were transcriptionally regulated in poplar or oak by all ECM interactions tested. Six of these (CAPE, CLE, GASA, nsLTP, PCY, and POE) were regulated by all ECM associations in both poplar and oak (Fig. 4 e and f). The highest proportions of genes regulated by all ECM interactions were found within the SSP families CAPE, KTI, and POE for poplar, and POE, PSY, and GASA for oak (Fig. 4 e and f). Of the ECM-responding SSPs, 17.3 % were similarly regulated by all the associations tested in poplar (18 genes) and oak (55 genes).

These results demonstrate that all 21 SSP symbiosis-related families, independently of their mode of action, are transcriptionally regulated by ECM associations in trees. They also showed that several SSP-encoding genes exhibited the same transcriptional response to all tested ECM interactions.

ECM-responsive PcCLE peptides stimulate the ECM association between *P. x canescens* and *L. bicolor*.

Among all SSP families, the CLE is one of the families whose functions and mode of action in symbiosis are best characterised (Nakagami et al., 2024). Short and therefore easily chemically synthesised, *in vitro* supplementation for pharmacological activity screens of CLE peptides in growth media identified several CLEs that regulate rhizobial and AM symbioses in herbaceous plants (Okamoto et al., 2013; Imin et al., 2018; Hastwell et al., 2019; Le Marquer et al., 2019). 34 CLE-encoding genes were regulated by ECM symbiosis in poplar (Fig. 5 a, Sup. Table S4). Therefore, they are good candidates for testing the effect of ECM-responsive peptides on ECM symbiosis in poplar.

**Fig. 5.**
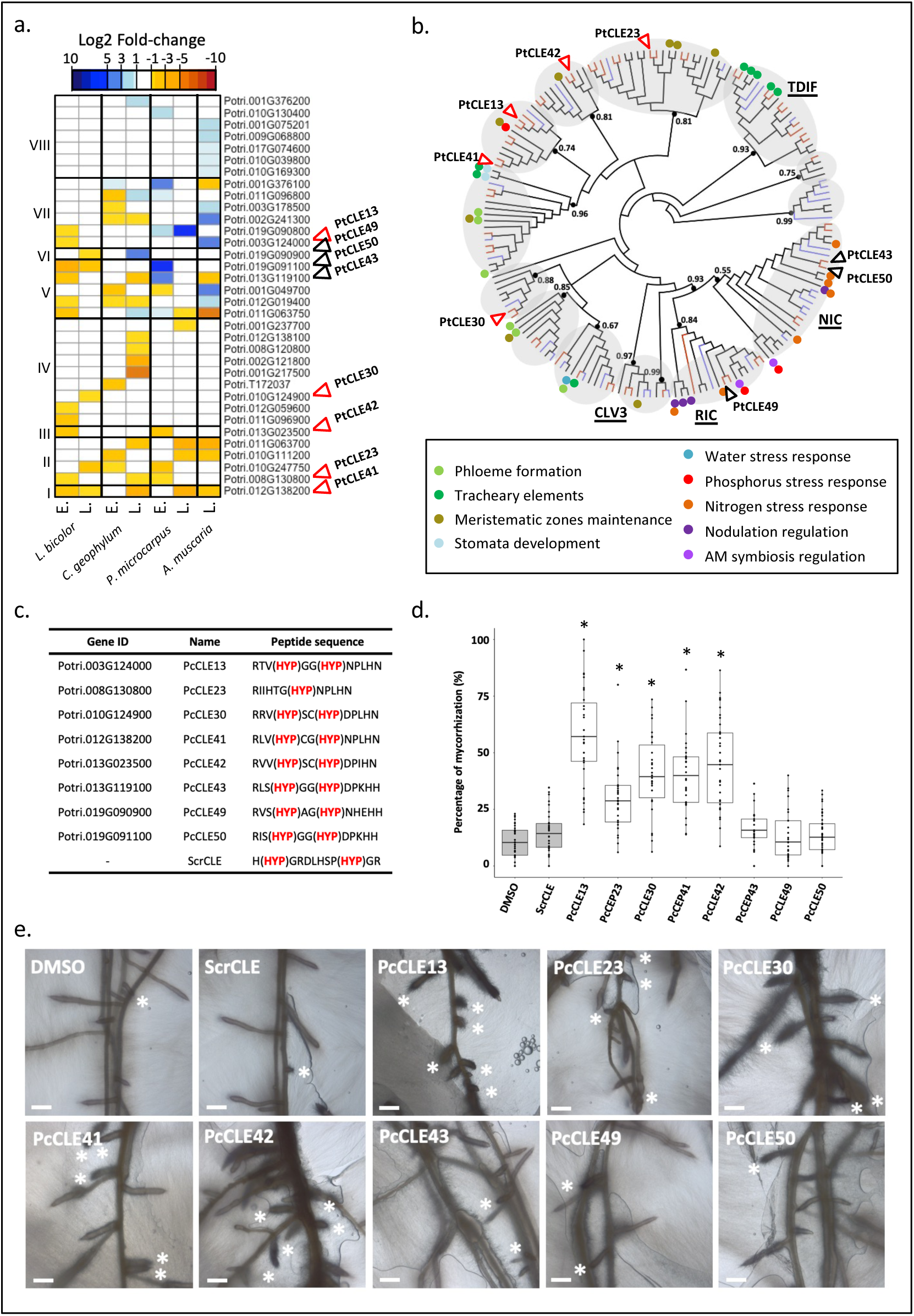
Five PcCLE peptides stimulated ectomycorrhizal associations between *P. × canescens* and *L. bicolor*. a. Heatmaps representing the clustering of CLE-encoding genes in poplar (*P. tremula **×** alba* for the interaction with *A. muscaria* and *P. **×** canescens* for all others), based on their pattern of relative expression in response to ectomycorrhizal associations with different fungi (log2 fold-change of the ratio of their expression in ectomycorrhiza compared to uncolonized roots). Conditions in which the regulation of a gene was not statistically significant (FDR > 0.05) are shown as not regulated (white cells). Roman numbers indicate different gene clusters based on the number of ectomycorrhizal associations that induce or repress the transcription of the SSP they comprise. Clusters numbers are the same as indicated in the legend of Figure 4. E. and L. are respectively an early and a late time point of sample collection after contact between the ectomycorrhizal fungi and the host tree (see Material and Method for details). b. Cladogram representations of the unrooted maximum-likelihood phylogeny of CLE pre-proteins present in the genomes of *A. thaliana* and *M. truncatula* (black branches of the cladograms), *P. trichocarpa* (red branches), and *Q. robur* (blue branches). The approximate likelihood ratio test (aLRT) support values are given for the major nodes of the trees and are marked with black points. Coloured points mark CLE peptides with known functions in herbaceous, the colour of the point indicates the associated function (Brand et al., 2000; Fiers et al., 2005; Hirakawa et al., 2010; Mortier et al., 2010; Araya et al., 2014; Liu et al., 2016; Gutiérrez-Alanís et al., 2017; Takahashi et al., 2018; Müller et al., 2019; Karlo et al., 2020; Kucukoglu et al., 2020; Mens et al., 2021; Zhang et al., 2022; Zhang et al., 2024). Localisations of the tested poplar CLEs (PcCLE13, 23, 30, 41, 42, 43, 49 or 50) in the tree are given (triangles). The red triangles mark the PcCLEs that stimulate the ectomycorrhizal association between *P. × canescens* and *L. bicolor*. c. Amino acid sequences of the *P. × canescens* PcCLEs (PcCLE13, 23, 30, 41, 42, 43, 49, or 50) and scrambled ScrCLE peptide (negative controls) tested on the ectomycorrhizal association between *P. × canescens* and *L. bicolor* and. (HYP) indicates the positions of hydroxylated prolines. d. Percentage of mycorrhization (number of lateral roots with mycorrhiza / total number of lateral roots) found between *P. × canescens* and *L. bicolor* co-cultivated on growth media supplemented with 1 µM of PcCLE peptide or with 1 µM of scrambled ScrCLE or DMSO (solvent) as negative controls. Asterisks mark treatments significantly (P-value ≤ 0.01) different from all controls (DMSO and ScrCLE), based on Kruskal-Wallis one-way analysis of variance and least significant difference (LSD) *post hoc* test corrected with the Bonferroni method, *n* = 25–53. Each dot represents a plant. e. Pictures of the root systems of *P. x canescens* co-cultivated with *L. bicolor* on growth media supplemented with 1 µM PcCLE peptides (PcCLE13, 23, 30, 41, 42, 43, 49, or 50) or with 1 µM scrambled ScrCLE or DMSO (solvent) as negative controls. White asterisks mark ectomycorrhizas. The scale bars represent 1 mm.

We selected eight CLE peptides whose encoding genes were transcriptionally induced by *L. bicolor* and were distributed across different expression clusters and phylogenetic clades (Fig.5 a and b). After verifying their CDS sequences by cloning, the corresponding peptides were synthesised with the addition of conserved proline hydroxylations known to be necessary for their functions in herbaceous (Fig. 5 c). After 2 weeks of co-culture on media supplemented with 1 µM of peptide (test conditions) or for negative controls with 1 µM of scrambled CLE (ScrCLE) and with the peptide solvent alone (DMSO), we found that PcCLE13, PcCLE23, PcCLE30, PcCLE41, and PcCLE42 increased significantly by two to three times the number of ECM roots formed between *P. × canescens* and *L. bicolor* (Sup. Fig. S7 a), and four to five times their percentage of mycorrhization (Fig. 5 d and e). PcCLE23 increased only the proportion of ECM roots. None of the five active peptides increased lateral root production, demonstrating that the observed increase in ECM root tips was not the result of the stimulation of lateral root formation (Sup. Fig. S7 b). However, all five active CLE peptides drastically repressed *P. × canescens* adventive root growth, decreasing their length by two to three times (Sup. Fig. S7 c and d).

Taken together, our results show that some of the poplar CLEs transcriptionally upregulated in *P. x canescens* - *L. bicolor* ECM roots stimulate *P. × canescens* ECM association with *L. bicolor*, increasing the number and proportion of ECM roots formed. Among the five active CLE peptides, PcCLE41 was transcriptionally induced in poplar by all ECM associations tested, and PcCLE23 and PcCLE42 were upregulated by at least two of them (Fig. 5 a). They may be involved in conserved processes of ECM symbiosis establishment or function. PcCLE30 was regulated by *L. bicolor* only, and PcCLE13 was oppositely regulated by different ECM associations in poplar. They may be involved in establishment processes specific to certain ECM interactions.

## Discussion

Plant SSPs are essential components of almost all processes necessary for plant growth and survival, including organogenesis, reproduction, adaptation to water and nutrient stresses, defence against pathogens, and rhizobial and AM symbioses (Tavormina et al., 2015; Zhang et al., 2025). In this study, we report for the first time the involvement of SSP of the CLE family in ECM symbiosis.

SSPs are produced from the translation of sORFs (<250 bp) or post-translational processing of longer preproteins (Matsubayashi, 2014; Tabata and Sawa, 2014; Tavormina et al., 2015). Classical gene prediction methods are biased against sORFs to limit the proportion of false discoveries, leading to their exclusion from genome annotations (Lease and Walker, 2006; Yang et al., 2011). In SSPs processed from preproteins, the family domain constituting the mature peptide, or family cysteine pattern, is conserved within members of the same SSP family; however, the rest of the sequence is not (Silverstein et al., 2007; Ohyama et al., 2008; Matsubayashi, 2014; Xie et al., 2022). This leads to their uneasy identification by global similarity searches (Oelkers et al., 2008; Ghorbani et al., 2015). Hence, SSPs are generally underrepresented in genomic annotations. To overcome this situation, most plant SSPs identification studies focus on large-scale *de novo* gene prediction inclusive of sORF (Lease and Walker, 2006; Hanada et al., 2007; Ohyama et al., 2008; Yang et al., 2011; Pan et al., 2012; Li et al., 2014; Ghorbani et al., 2015; Hu et al., 2022) or on the identification of the members of individual SSP families by motif similarity searches (Delay et al., 2013; Takata et al., 2013; Guo et al., 2015; Cao et al., 2015; Goad et al., 2017; Hastwell et al., 2017; Wang et al., 2021b; Wei et al., 2023). In 2017, De Bang et al. (2017) combined both approaches. The power of detection of *de novo* gene prediction associated with the precision of motif similarity searches by HMM and a careful manual curation identified the members of 46 SSP families of interest in *M. truncatula*. We used the same strategy to generate for two ECM tree species, *P. trichocarpa* and *Q. robur,* the gene repertoire of 21 SSP families with known members involved in AM or rhizobial symbioses or transcriptionally regulated by these associations in herbaceous plants and present in trees (Sup. Tab. S1). We identified 607 SSPs in *P. trichocarpa* and 446 SSPs in *Q. robur*. Among these, 23% of the poplar and all the oak sequences were new. We found that all Poplar and Oak SSPs shared the sequence characteristics of their families, suggesting the conservation of modes of action and functions of SSP families between herbaceous plants and trees. The high and consistent recovery rates obtained for all 21 families, independently of their categories (PTM, FP, Cys-rich, sORF, or others), along with the identification of novel members for several SSP families in both test species (*A. thaliana* and *M. truncatula),* strongly suggest that the pipeline and captured the integrity of the selected 21 SSP families across the four species. The identification and correct family assignment of four published SSPs of *A. thaliana*, *M. truncatula,* and *P. trichocarpa,* despite lacking conserved family domains, support this claim.

*De novo* gene prediction from genome yielded little new genes (9,2 %), except for *Q. robur* (41,7 %). These included *Q. robur* closest homologous of well-characterised peptides, such as CLV3 or IDA (Sup. Table S3) justifying *de novo* gene prediction. The large difference in the outcome of SSP-targeted *de novo* gene prediction between *Q. robur* and the other species is concordant with a lower depth of SSP annotation of *Q. robur*. It can be explained by the recency of the genome compilation of *Q. robur* (Plomion et al., 2018) which has not, contrary to those of *A. thaliana* (Lamesch et al., 2012), *M. truncatula* (Tang et al., 2014) and P*. trichocarpa* (Tuskan et al., 2006), undergone many rounds of annotation improvements.

To identify plant peptides with roles in ECM symbiosis among the 21 symbiosis-related SSP families, we examined their transcriptional responses to various ECM associations in poplar (*P. × canescens* and *P. tremula × tremuloides*) and oak (*Q. robur*). 48.6% of poplar SSPs and 28.5% of oak SSPs, were regulated by at least one ECM interaction. These included members of all 21 SSP families in poplar and of 16 in oak. High percentages of regulated SSPs were also reported in these families for AM and rhizobial symbioses (de Bang et al., 2017). In selecting symbiosis-related SSP families, we intentionally focused on those likely to be regulated by symbiotic associations. This bias probably contributed to the high proportion of regulated SSP families and encoding genes. Fewer SSP-encoding genes and families regulated by ECM symbiosis were found in oak than in poplar. This difference is likely a consequence of the limited diversity and number of ECM interactions examined in oak (three *Tuber* species) compared to poplar (four species from four different families and two phyla). This narrower range of ECM interactions probably led to the identification of fewer SSP specific to single ECM associations, resulting in a lower total number of regulated SSPs and SSP families identified in oak. Diversifying the range of ECM associations would likely increase the number of ECM-regulated SSPs in both trees. These results suggest that likewise in AM and rhizobial symbiosis (de Bang et al., 2017), many SSP families are involved in ECM symbiosis.

No bias was observed in the ECM regulation of any SSP family or mode of action (signalling, antimicrobial, and peptidase inhibitor peptides), implying that SSPs participating in the ECM process support a diversity of functions. This diversity already appeared in the study by Plett et al. (2017), who also identified signalling, antimicrobial, and peptidase inhibitor peptides in *P. × canescens* root transcripts differentially regulated by *L. bicolor*. Only 34 regulated SSPs were common to both studies, accounting for 3.5 % of the total SSPs regulated in both studies. Such large differences can be attributed to the use of different prediction approaches (prediction of SSPs by the size of their ORFs and the presence of a signal peptide in the N-terminus versus search for specific families integrating *de novo* gene prediction).

Because the ability to form ECM symbiosis evolved convergently multiple times in both trees and fungi (Tedersoo et al., 2010; Tedersoo and Brundrett, 2017), it is possible that no universal ECM molecular pathway exists and that ECM interactions differ in the ways they are established and regulated across tree species. Alternatively, isolated molecular tools may have been co-opted independently by different tree species to allow or control fungal colonisation. To address this, it is essential to compare ECM models associating different trees and fungal species. Phylogenetic analysis indicated that ECM symbiosis evolved independently in *Populus* and *Quercus* (Wang and Qiu, 2006). When comparing the regulation of SSPs in poplar and oak, *oak* exhibited more SSP families and SSP-encoding genes regulated by all interactions. This difference is likely due to the closer phylogenetic relationship between the fungal species tested in oak, which may lead to a higher number of common molecular mechanisms and, as a result, *to* a greater number of conserved SSP families regulated by all interactions. We identified 13 SSP families with members regulated by all tested ECM interactions in oak or poplar. Among them, the identity of the families with the highest proportion of genes regulated by all ECM interactions varied between poplar and oak. However, six of these families (CAPE, CLE, GASA, nsLTP, PCY, and POE) were shared between the two tree species, suggesting that they may harbour common determinants of ECM symbiosis. The CLE, POE and PCY families encode signalling-SSPs (Luo et al., 2018; Kim et al., 2021; Zhang et al., 2025), while CAPE, GASA, and nsLTP peptides are associated with antimicrobial functions without knowledge of their mode of action (Breen et al., 2017; Fleury et al., 2019; Gao et al., 2022; Han et al., 2023; Nahirñak et al., 2024). No common regulation patterns of these families emerged between poplar and oak. An in-depth analysis of their expression patterns showed that within these families, most SSP-encoding genes regulated by all ECM interactions in poplar or oak followed different expression patterns in response to different interactions. Given the diversity of their regulation, and in the absence of evidence of their roles in ECM symbiosis, it is currently impossible to conclude whether this large number of SSP families advocate in favour of a common ECM pathway, are co-opted molecular tools, or only respond to these interactions. Nevertheless, several SSP-encoding genes (18 in poplar and 55 in oak) were similarly regulated by all the tested associations (clusters I and XI, Sup. Table S4 and clusters II and X, Sup. Table S5). They may be involved in a general process of ECM symbiosis establishment and function in these tree species. At the opposite, the SSPs regulated by only one interaction may play a specific role in the establishment of this particular interaction.

In herbaceous plants, some CLEs vehiculate satiety signals repressing both at the local and systemic levels rhizobial and AM symbioses in response to phosphorus and nitrogen availability or to prior rhizobial and AM associations (autoregulation signals) (Okamoto et al., 2008; Mortier et al., 2010; Reid et al., 2011; Nishida et al., 2016; Müller et al., 2019; Karlo et al., 2020; Mens et al., 2021; Wang et al., 2021a; Wulf et al., 2024). CLE peptides are also crucial for meristematic maintenance and vascular tissue differentiation (Hirakawa et al., 2010; Zhu et al., 2020b; Zhang et al., 2024). Leveraging these developmental functions, several parasitic and symbiotic organisms co-opted CLE-encoding genes to ease their accommodation during root colonisation. For example, CLE-encoding genes are present in the genomes of certain glomeromycete species and are expressed during their interactions with the host plants (Le Marquer et al., 2019). They enhance root colonisation by promoting lateral root emergence and development. Similarly, certain nematodes express CLE genes that stimulate vascular development, aiding the formation of feeding sites in root tissues (Replogle et al., 2011; Wang et al., 2011; Chen et al., 2015; Guo et al., 2017). As one of the six SSP families regulated by all ECM interactions in poplar and oak, CLE peptides were ideal candidates for testing the association between the transcriptional response of SSPs to ECM symbiosis and their role in the establishment or regulation of these associations. We found that five PcCLEs (PcCLE13, PcCLE23, PcCLE30, PcCLE41, and PcCLE42) increased the formation of ECM roots between *P. x canescens* and *L. bicolor*, marking the first evidence of CLE involvement in ECM symbiosis. Strikingly, while CLE peptides are known repressors of AM and rhizobial symbiosis in herbaceous plants, they stimulate the ECM association of poplar and *L. bicolor*. This demonstrates that although PcCLEs are induced by ECM associations, they are not involved in ECM autoregulation. Rather, they may participate in the establishment of ECM associations or formation of ECM roots.

All known CLE-encoding genes that regulate AM and rhizobial interactions belong to two affiliated phylogenetic clades (Fig. 5 b) : RIC (RHIZOBIA-INDUCED CLE) and NIC (NITRATE-INDUCED CLE) (Reid et al., 2011; Hastwell et al., 2014; Goad et al., 2017; Hastwell et al., 2017; Mens et al., 2021). None of the three tested PcCLEs belonging to these two clades affected ECM root formation (Fig. 5 b), suggesting that CLE function in ECM symbiosis is different from that of RIC and NIC CLEs in AM and rhizobial associations. Instead, all active PcCLEs were members of clades associated with CLE peptides that regulate meristematic activities in herbaceous and poplar (Fiers et al., 2005; Hirakawa et al., 2010; Liu et al., 2016; Gutiérrez-Alanís et al., 2017; Zhang et al., 2022; Zhang et al., 2024). Young lateral roots are the sites of ECM symbiosis. An increase in lateral root formation might explain the observed ECM stimulation. However, none of the active PcCLEs increased the number of lateral roots. This shows that, contrary to the effects of glomeromycete CLEs, the stimulation of ECM association is not the result of enhanced lateral root formation. We found that a common feature of active PcCLEs was their inhibitory effect on poplar adventitious root growth. ECM root organogenesis involves root apical meristem arrest and full differentiation of colonised roots (Felten et al., 2009; Vayssières et al., 2015). Thus, it is possible that the expression of active PcCLEs controls the lateral root meristem differentiation necessary for ECM root formation. ECM fungi are known to manipulate host pathways (Daguerre et al., 2020; Kang et al., 2020; Plett et al., 2020b; Wong-Bajracharya et al., 2022). Whether the expression of active PcCLEs is part of the plant ECM pathway or are highjacked by ECM fungi to facilitate colonisation remains to be determined. Among the five active CLEs, only PcCLE41 was transcriptionally induced by all ECM associations tested. PcCLE23 and PcCLE42 were upregulated by two ECM associations, PcCLE30 was upregulated by *L. bicolor* only, and PcCLE13 was oppositely regulated by different ECM associations. These differences in expression patterns suggest that both common and specific ECM association processes coexist in a single tree species.

Although only 23.5% of the CLE-encoding genes regulated during ECM symbiosis were examined in our study, 62.5% of the selected genes interfered with ECM symbiosis. CLE peptides accounted for only 8% of ECM-regulated SSPs, and their effects were analysed exclusively for ECM root formation. We did not investigate other phenotypes, such as the structure of the ECM roots or their active functioning, leaving a substantial genetic and functional gap to explore. Therefore, it is probable that numerous other ECM-regulated SSPs may play a role in the establishment or functioning of ECM symbiosis. Although this study provides significant insights into the role of SSPs in ECM symbiosis, further research is necessary to elucidate their precise functions, modes of action, and reveal the molecular mechanisms involved in their recruitment.

## Methods

### *In-silico* identification of SSP-encoding genes (Identification pipeline)

The members of 21 SSP families were searched in the genome assemblies and predicted proteomes of *Arabidopsis thaliana* (TAIR10; Lamesch et al., 2012), *Medicago truncatula* (Mt4.0v1; Tang et al., 2014), *P. trichocarpa* (v3.1; Tuskan et al., 2006), and *Q. robur* (PMN1; Plomion et al., 2018). Each predicted proteome was mined with hmmscan (HMMER package) using the Hidden Markov Model (HMM) profiles of each SSP family defined by de Bang et al. (2017). To enhance our set of reference sequences, if the hmmscan result contained at least five sequences, we attempted to discover new motifs using MEME (MEME suite; Bailey et al., 2015). We used the new motifs generated to perform a MAST scan (MEME suite) of the predicted proteome with default parameters. We used these reference sequences as queries for similarity searches with tBLASTn on the genome assembly. Scaffolds containing at least one match to a reference sequence (e-value < 1e^-5^, identity > 30 % and coverage > 60 %) were extracted and used for *de novo* gene prediction with Augustus (Keller et al., 2011) (species parameter set to *A. thaliana*). *De novo* predicted proteins containing an SSP HMM profile and/or having a BLASTp hit with one reference sequence (e-value < 1e^-5^, identity > 30 % and coverage > 50 %) were selected. We defined the pool of predicted SSPs as the non-redundant combination of the reference sequences and their homologous sequences identified from *de novo* gene prediction. To achieve non-redundancy, we compared this new set of proteins with the predicted proteomes and eliminated duplicates.

For each SSP family, manual curation was achieved using CLANS (Frickey and Lupas, 2004) to perform the all by all BLASTP of the full amino acid sequences of the predicted SSPs of the four species, and their clustering according to their p-value (cut-off 10e^-14^). True cluster(s) were identified based on the presence of known sequences within and at proximity, and on the density of connections within. Sequences that formed outgroups, pairs, or singletons with no other connections to the true clusters were filtered out and compared by amino acid alignment with the remaining predicted sequences. Alignments were performed with MAFFT version 7 (Katoh and Standley, 2013) using the iterative refinement strategy E-INSI (for the DVL and POE families), G-INSI (for the TAX family), or L-INSI (for all other families). This comparison allowed the identification of putative pseudogenes with interrupted open reading frames and prediction errors (false positives). Interrupted sequences were retained, and false positives removed.

### Clustering, Phylogenetic analysis, SSP domain identification and localisation predictions

Pairwise similarity clustering was performed by CLANS using full amino acid sequences (p-value cut-off 10e-^14^) of all identified peptides (Frickey and Lupas, 2004). Phylogenies were produced from the alignments (performed as described in the previous paragraph) of the full amino acid sequences. Sequences with interrupted ORF were included in the analysis to increase resolution (Jiang et al., 2014). The sequence alignments were not trimmed. Maximum-likelihood phylogenetic trees were generated with ATGC PhyML using the approximate likelihood ratio test (aLRT) to calculate the statistical support values of each embranchment (Guindon et al., 2010). The embedded automatic model selection (SMS) of PhyML was used to determine the amino acid substitution model best fitted to the phylogenetic analysis of each SSP family (Lefort et al., 2017). Trees were visualised using FIGTREE (http://tree.bio.ed.ac.uk/software/figtree/).

MEME (Bailey and Elkan, 1994) was used with a constraint of one to six conserved motifs and no limited number of motif repetitions to assess the number of SSP family conserved domains in the sequences of the SSP preproteins. LOGO representations of the conserved domains were generated using WEBLOGO (Crooks et al., 2004). Subcellular localisations were predicted using TargetP version 2 (Armenteros et al., 2019), SignalP version 6 (Teufel et al., 2022), and DeepLoc version 2 (Thumuluri et al., 2022), specifying the plant origin of the sequences and using default settings.

### Transcriptomes and differential gene expression analysis

RNA-Seq datasets of poplar root interaction with *Laccaria bicolor (S238N)*, *Cenococcum geophillum (1.58), Pisolithus microcarpus* (*441)* and *Amanita muscaria* (*MEII)* and of oak root interaction with *Tuber melanosporum, Tuber aestivum and Tuber magantum* were retrieved from the Short Read Archive (SRA) and the GEO and ENA databases (Sup. Table S6). Details of their production are published in Basso et al., (2020), de Freitas Pereira et al., (2018), Marqués-Gálvez et al., (2025) and Kohler et al., (2015) for poplar, and in Plomion et al., (2018) for oak. For each dataset, raw reads were paired, trimmed for quality and sequencing adapters, and mapped against *Populus trichocarpa* v3.1 (https://phytozome-next.jgi.doe.gov/info/Ptrichocarpa_v3_1) or *Q. robur* haplome (Qrob_PM1N_CDS_nt_20161004, https://www.oakgenome.fr/index8568.html?page_id=587) references transcripts to which the gene sequences newly identified in this study were added, using CLC Genomics Workbench v21 (Qiagen). The following CLC parameters were used for mapping: minimum length fraction 0.9, minimum similarity fraction 0.9, mismatch cost 2, insertion cost 3, deletion cost 3, and maximum number of hits for a read 10. Unique reads counts were exported for further analyses. Normalised expression values were generated and differential transcription levels of the reads calculated with DESeq2 (v1.44.0) (Love et al., 2014). The transcriptome of each plant-fungal interaction was compared to the transcriptome of non-inoculated plants cultivated in the same conditions. An early stage and a late stage of ectomycorrhizal root development were used for poplar interactions with *L. bicolor* (1 and 2 weeks), *C. geophylum* (3 and 8.5 weeks) and *P. microcarpus* (1 and 4 weeks), except for *A. muscaria* (only late stage, 8 weeks). Condition-specific differentially expressed genes with a false discovery rate (FDR) < 0.05 and a log2 fold-change (Log2FC) ≥ |1| were identified for each pairwise condition. Log2FC of relative expression (expression in ECM compared to uncolonized roots) was used to generate the heatmaps (R package pheatmap, RRID: SCR_016418). Clustering was performed based on the number of ECM associations that induced or repressed statistically significantly the transcription of each gene. Bubble plots were produced using the numbers and proportions of SSP-encoding gene that were statistically significantly regulated by one, several or all the ECM fungi tested for each SSP family (R package ggplot2, RRID: SCR_014601).

### Peptides synthesis

Total RNA was extracted from grounded frozen *P. × canescens – L. bicolor* ECM roots using the RNeasy Plant Mini Kit (Qiagen, Hilden, Germany) and DNase treated with the Turbo DNae-free Kit (Thermo Fisher Scientific, Waltham, AS, US). cDNAs were synthesized from 1 µg of total DNAse-free RNA using the SuperScript III First-Strand Synthesis Kit (Invitrogen, Waltham, AS, US) with oligo(dT)_20_ primers. The CDSs of interest were verified by PCR amplification from 1 µL of cDNA using the Phusion High fidelity DNA polymerase (Thermo Fisher Scientific, Waltham, AS, US) (primers and melting temperatures in Sup. Table S7), cloning into pGEMTeasy vectors (Promega, Madison, WI, US) and sanger sequencing. The amino acid sequences encoded by these CDSs were deduced using Expasy Translate (Gasteiger et al., 2003). The CLE peptides and scrambled counterpart (ScrCLE) were synthesised by Genecust (Boynes, France) with proline hydroxylations on proline 4 and 7.

### Biological material and growth conditions

Clones of *Populus × canescens* (*Populus tremula × Populus alba* line INRA 717-1B4) were maintained *in vitro* by micropropagation in glass tubes on 1,2 % agar half-strength Murashige and Skoog (MS/2) medium (Murashige and Skoog, 1962) at 20°C under a 16h photoperiod at a light intensity of 150 µmol/(m2 . s) (OSRAM Fluorescent tubes 50/50 Fluora/Cool white). Dikaryotic vegetative mycelium of *L. bicolor* S238N (Maire P. D Orton) was maintained by *in vitro* subculture on 1,2 % agar modified Pachlewski medium (P5) (Deveau et al., 2015) at 25°C in the dark.

For cloning, total RNA was obtained from *P. × canescens – L. bicolor* ectomycorrhiza produced by *in vitro* co-cultivation according to the sandwich method (Felten et al., 2009). After two weeks of co-cultivation, ectomycorrhiza were collected, flash-frozen in liquid nitrogen, and conserved at −80°C.

For the CLE activity assay, cuttings of *P. × canescens* grown 20 days between cellophane membranes on 0.8 % agar MS/2 media were transferred onto cellophane membranes on 0.8 % agar MS/2 media supplemented with 1 µM of synthetic peptides for two weeks of pre-treatment. Following pre-treatment, poplars were co-cultivated two weeks with *L. bicolor* on fresh 0.8 % agar MS/2 media supplemented with 1 µM of the same synthetic peptide. Peptides were dissolved in dimethyl sulfoxide (DMSO) and added to the growth medium before pouring the plates. Scrambled peptides or addition of the same volume of DMSO without peptides served as negative controls.

### Image acquisition and phenotypic analysis

After two weeks of co-cultivation, ECM colonisation was assessed expressing the rate of mycorrhization as the percentage of ectomycorrhiza within the total number of lateral roots. Root systems were photographed under a Discovery V.8 stereomicroscope (Zeiss). The lengths of the adventive roots were measured on scans of the co-cultivation plates obtained with an Epson Perfection V700 PHOTO scanner (EPSON, Suwa, Japan), using the image processing package FIJI (https://fiji.sc). 25 to 53 plants per conditions were observed. Differences between treatments and controls (DMSO and ScrCLE) were tested for each root parameter (ECM root number, total lateral root number, and adventive root length) using the Kruskal-Wallis one-way analysis of variance and post-hoc Fisher’s LSD corrected with the Bonferroni method (R package agricolae).

## Data and code availability

The scripts for the SSP identification pipeline are available at https://forge.inrae.fr/iam/mycor_plant_ssps. Transcriptomes were retrieved from previous studies; the accession numbers are summarized in Supplemental Table S6.

## Acknowledgments

This work was funded by the Laboratory of Excellence Advanced Research on the Biology of Tree and Forest Ecosystems (ARBRE) (ANR-11-LABX-0002-01). The authors acknowledge Dr. Félix Fracchia for his advices on redaction and figures.

## Author contributions

C. B. and F. M. obtained funding and conceptualized the study; C. B., E. M., F. M., A. K., and C.V.-F. designed the experiments; C. B. and E. M. designed the bioinformatic pipeline; E. M. developed and operated the bioinformatic pipeline; C. B. performed the manual curation of the pipeline output; A. K. and C. B. performed the transcriptional analysis; C. B., E. d. S. M., A. H. and A. C. verified the CDS sequences by cloning; C. B. and A. C. tested *in vitro* the activity of the CLE peptides; C. B. performed and analysed all other experiments. The original manuscript was drafted by C. B; E. M., F. M., A. K. and C.V-F reviewed and edited the manuscript.

## supplemental figures

**Sup. Fig. S1.**
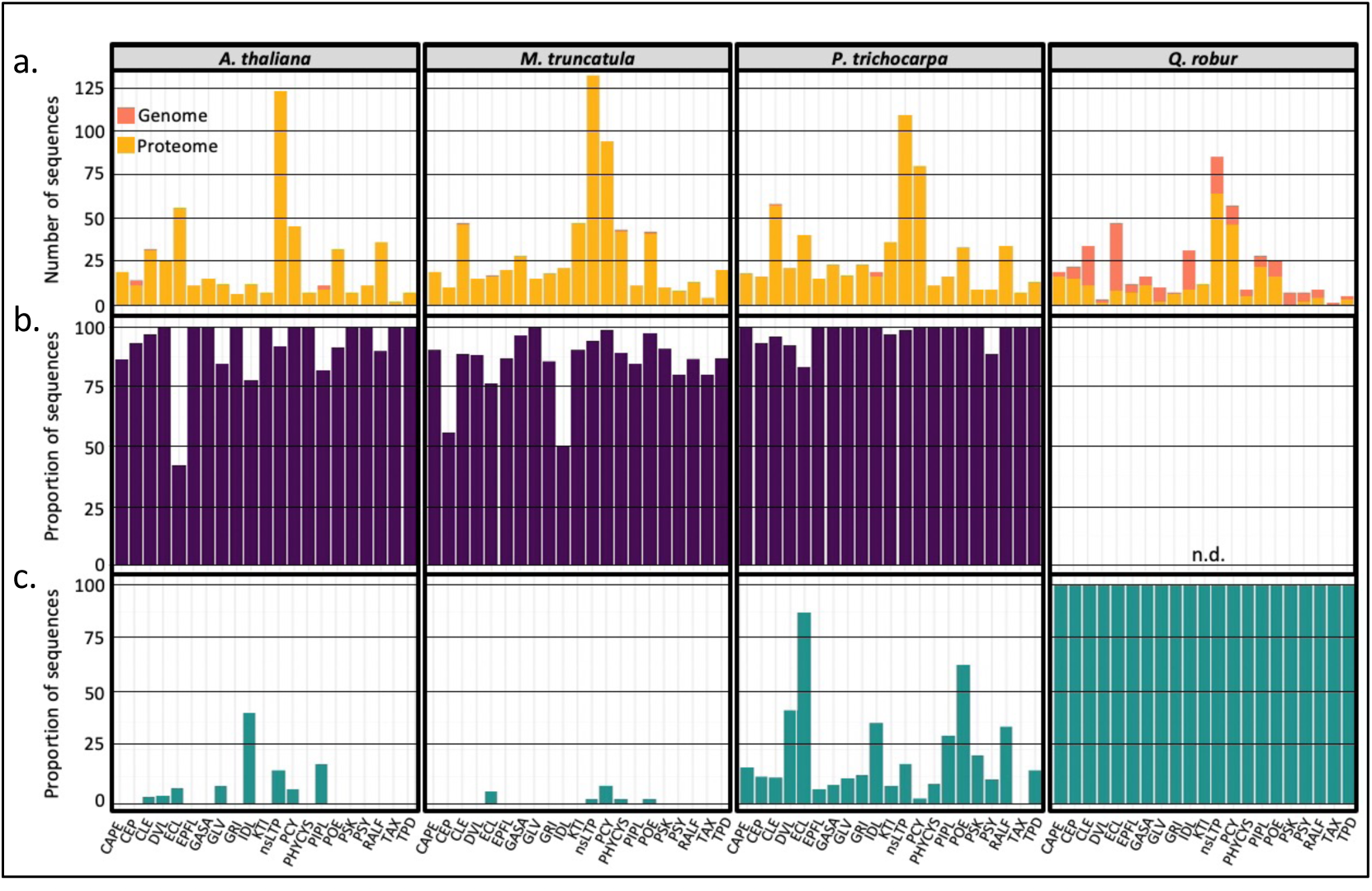
SSP members of the 21 symbiosis-related SSP families in *A. thaliana*, *M. truncatula*, *P. tricocharp*a and *Q. robur.* a. Number of SSP sequences of each of the 21 symbiosis-related SSP families identified by the Identification pipeline from the predicted proteomes and genomes of *A. thaliana*, *M. truncatula*, *P. tricocharpa* and *Q. robur*. b. Percentage of published SSP recovered by the Identification pipeline for each of the 21 symbiosis-related SSP families from *A. thaliana, M. truncatula, P. tricocharpa* and *Q. robur*. c. Percentage of novel SSPs identified by the Identification pipeline for each of the 21 symbiosis-related SSP families in *A. thaliana*, *M. truncatula*, *P. tricocharpa* and *Q. robur*.

**Sup. Fig. S2.**
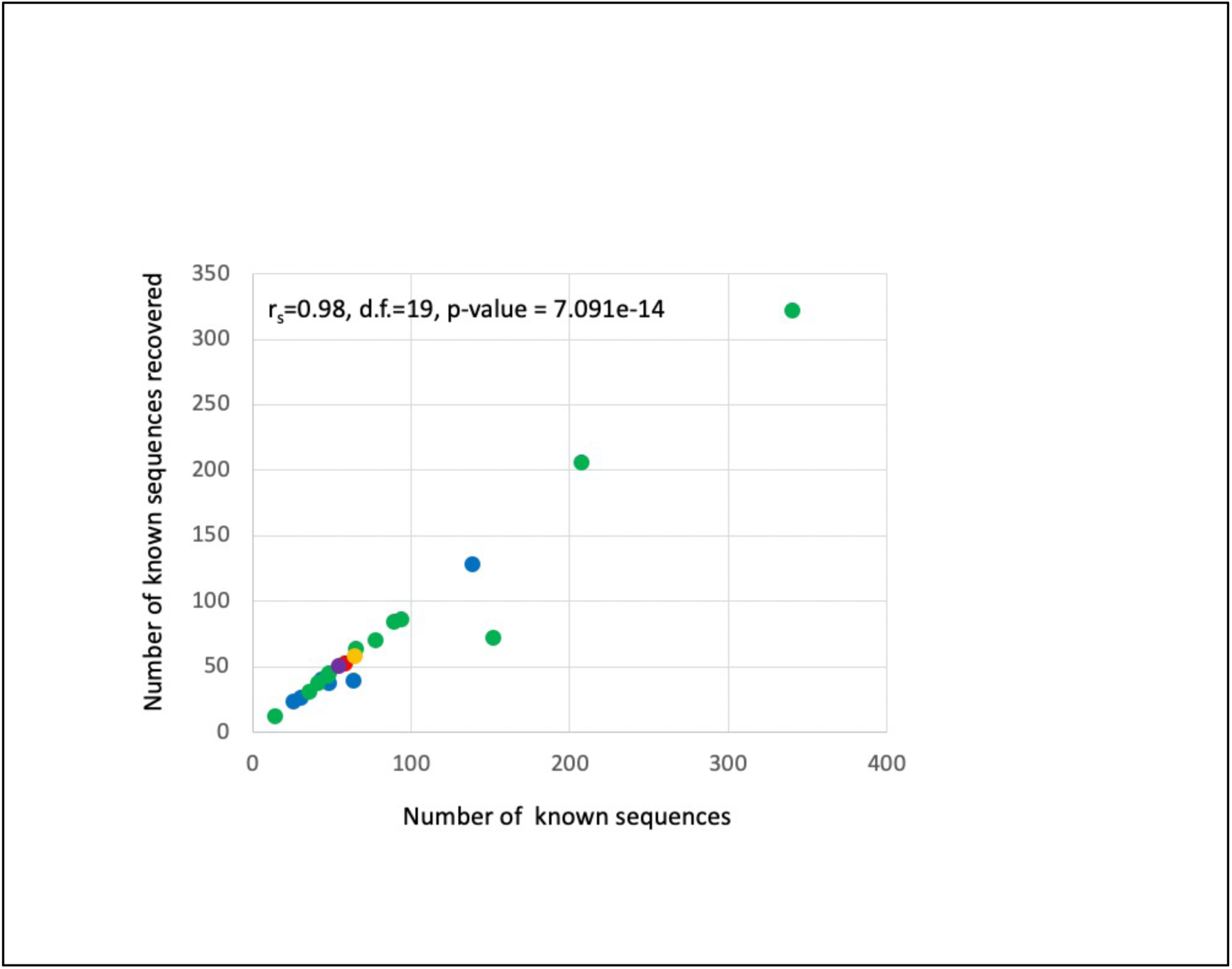
Verification of Identification biases associated with the SSP family or family categories in our pipeline. Recovery correlation plot displaying the number of known sequences recovered by the Identification pipeline as a function of the number of known (published) sequences in each of the 21 SSP families. Each point represents an SSP family. Different colours represent different family categories: green for cysteine-rich families (C-rich), blue for Post-Translational Modified peptide families (PTMs), red for the families of peptides produced from functional precursors (FP), purple for the families of peptides produced from small ORFs (sORFs), and yellow for those not entering into any of the previous categories (Other). The non-parametric Spearman rank-order correlation coefficient (r_S_) was used to verify the statistical significance of the correlation between the number of recovered sequences and the number of known (published) sequences.

**Sup. Fig. S3.**
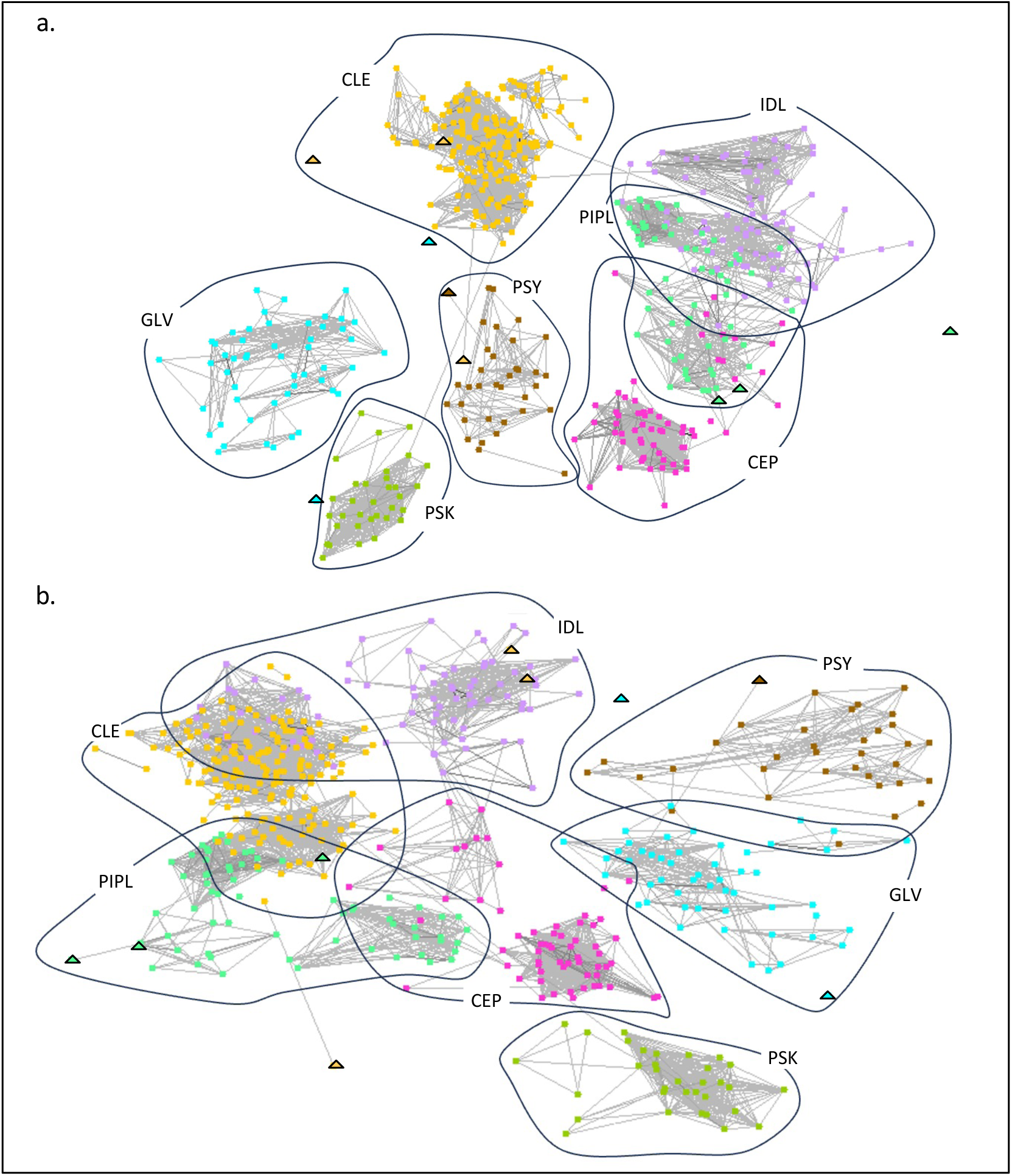
SSP members of the seven post-translationally modified SSP families of interest were well discriminated. Two-dimensional views of pairwise similarity three-dimensional clustering diagrams produced by CLANS (Frickey and Lupas, 2004) from the full amino acid sequences encoded by the SSP-encoding genes identified for the seven post-translationally modified SSP families (CEP, CLE, GLV, IDL, PIPL, PSK, and PSY). Two different angles of view (a and b) of the same three-dimensional clustering are display to demonstrate good discrimination of the SSP families. Each peptide is represented by a square whose colour indicates the SSP family to which it was affiliated. Peptides are connected by a grey line when the *P-*value of their reciprocal BLASTP hits is P ≤ 10^-10^. The attractive forces between peptides used for clustering were proportional to the negative logarithm of the peptides reciprocal BLASTP *P-values*. Triangles mark SSP sequences clustering at the outskirts of their family (the CLEs Medtr1g106920, Medtr7g093050, and Medtr7g084110; the PIPLs AT3G06090, AT3G13370, and prot24978; the GLVs AT5G51451 and Potri.010G147100; and the PSY AT3G49270).

**Sup. Fig. S4.**
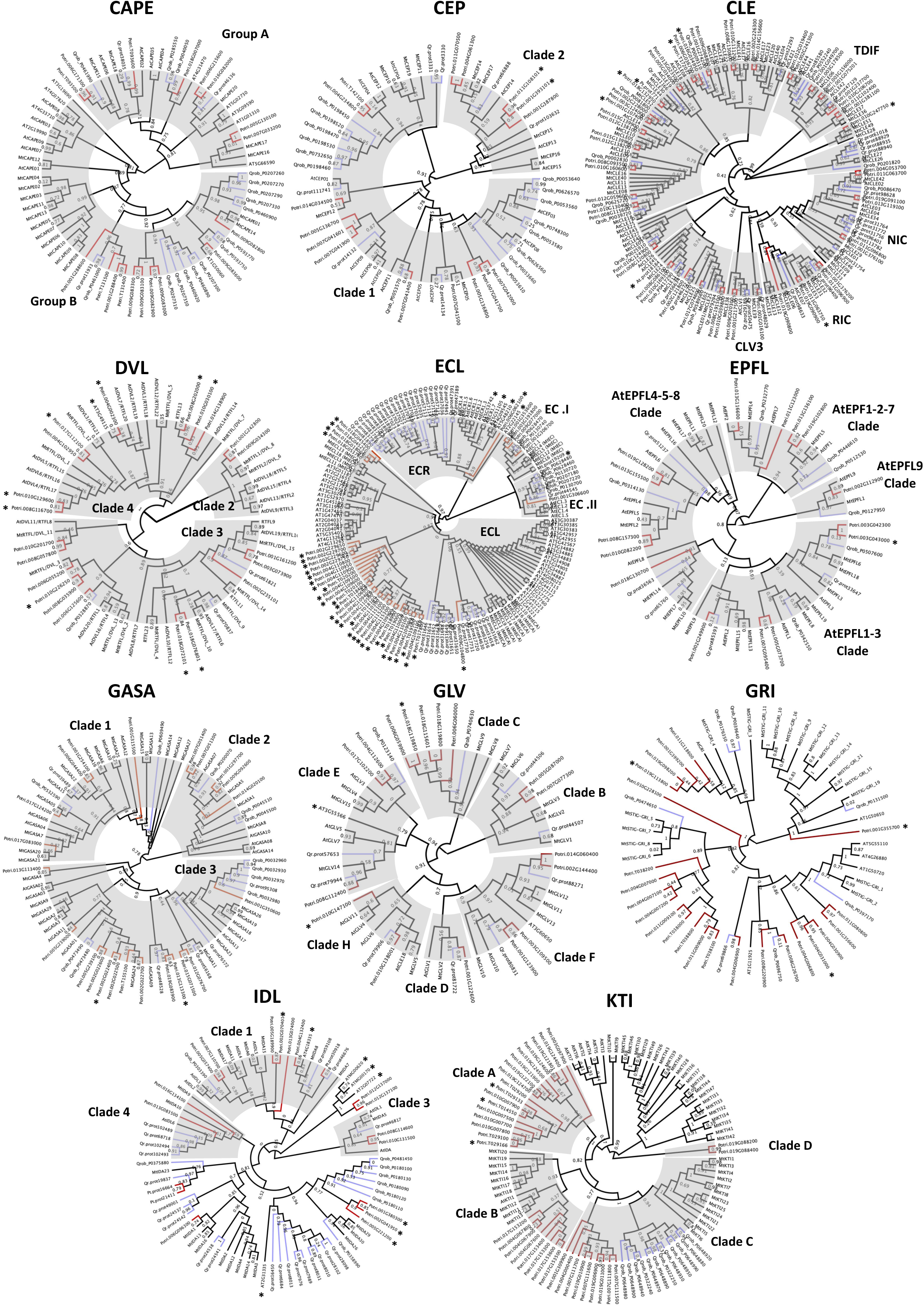
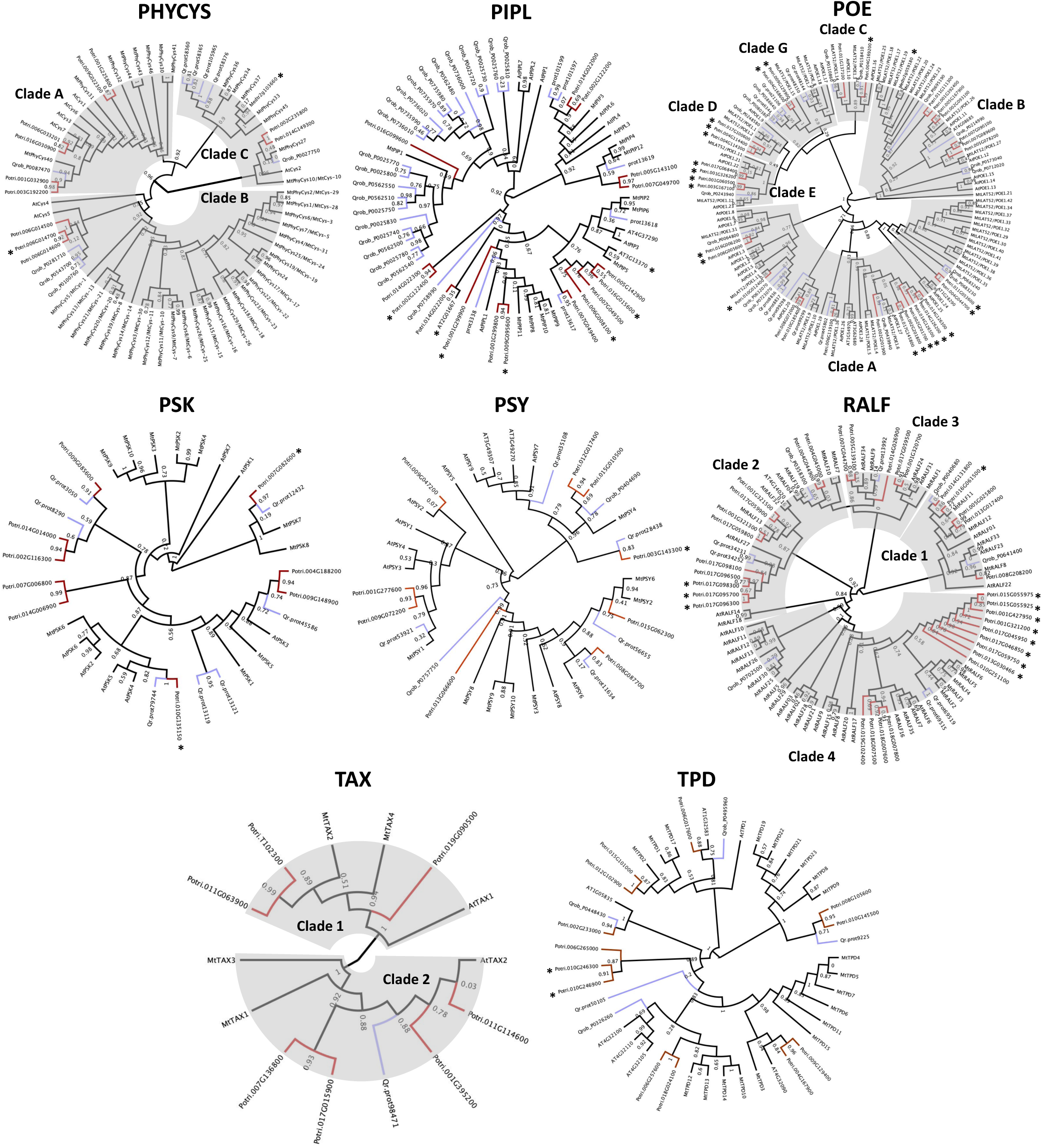
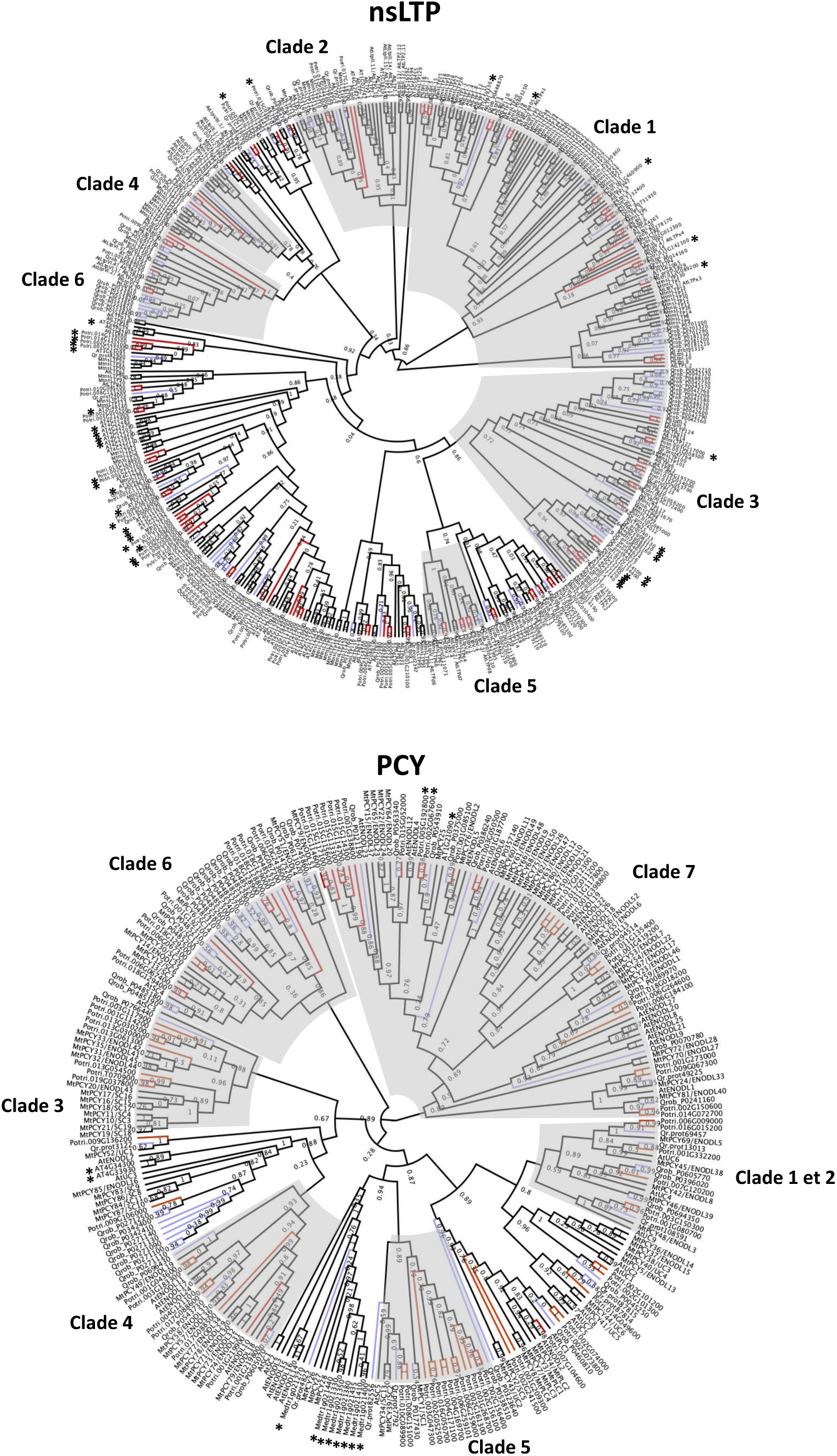
Phylogenies of the 21 selected SSP families. Individual cladogram representations of the unrooted maximum-likelihood phylogenies of the 21 selected families. Each of the 21 phylogenies were produced from the alignment of the full amino-acid sequences encoded by the ORFs of all genes identified by the prediction pipeline in the genomes of the herbaceous *A. thaliana* and *M. truncatula* (black branches of the cladograms) and of the ectomycorrhizal trees *P. trichocarpa* (red branches) and *Q. robur* (blue branches). For sequences of *A. thaliana* and *M. truncatula,* names consist of the species code (first letter of the genus and first letter of the species name) and gene names as published when existing; identifiers of the genes are given otherwise. Approximate likelihood ratio test (aLRT) support values were provided for all nodes of the cladogram. The clades described in the literature are marked by grey zones. Missing clades were due to unrecovered sequences or absence of the organisms in which these clades were present in our study. The name or number identifying each of these clades is provided as published in previous studies (CAPE (Wang et al., 2023); CEP (Delay, 2015); CLE (Zhang et al., 2020); DVL (Guo et al., 2015); ECL for EC clade see (Wang et al., 2021b), for ECR and ECL clades (no published phylogenies); EPFL (Takata et al., 2013); GASA (Wu et al., 2022); GLV (Fang et al., 2021); GRI (no published phylogeny); IDL (Guo et al., 2021); KTI (Ma et al., 2011); nsLTP (Wei et al., 2023); PCY (Luo et al., 2018); PHYCYS (He et al., 2021); PIPL (Hou et al., 2014); POE (Qian et al., 2020); PSK (Di et al., 2022); RALF (Campbell and Turner, 2017); TAX (Gholami, 2013); PSY and TPD no published phylogenies). Asterisks mark the novel sequences identified in *A. thaliana*, *M. truncatula* and *P. trichocarpa*. All *Q. robur* sequences were novel.

**Sup. Fig. S5.**
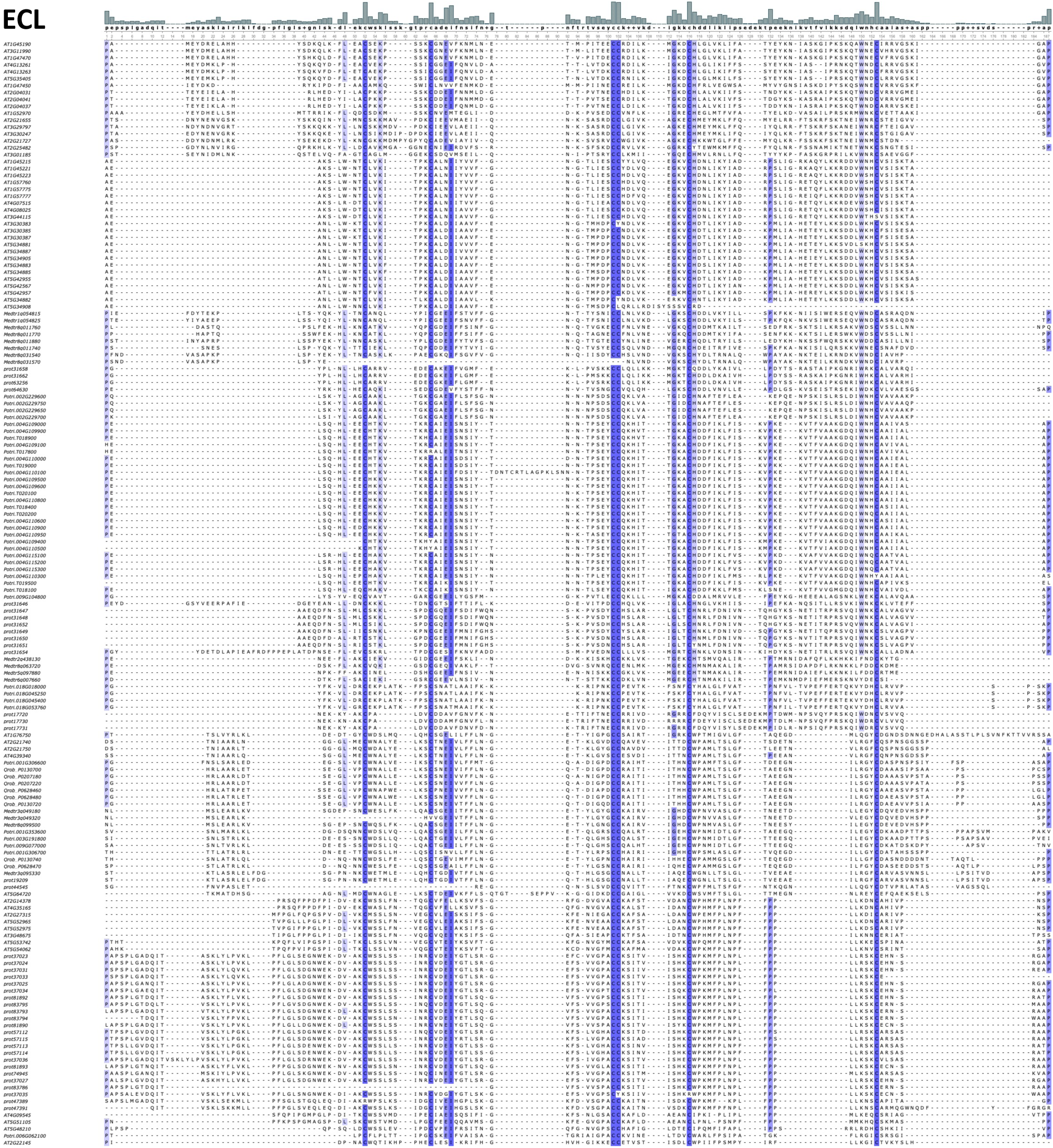
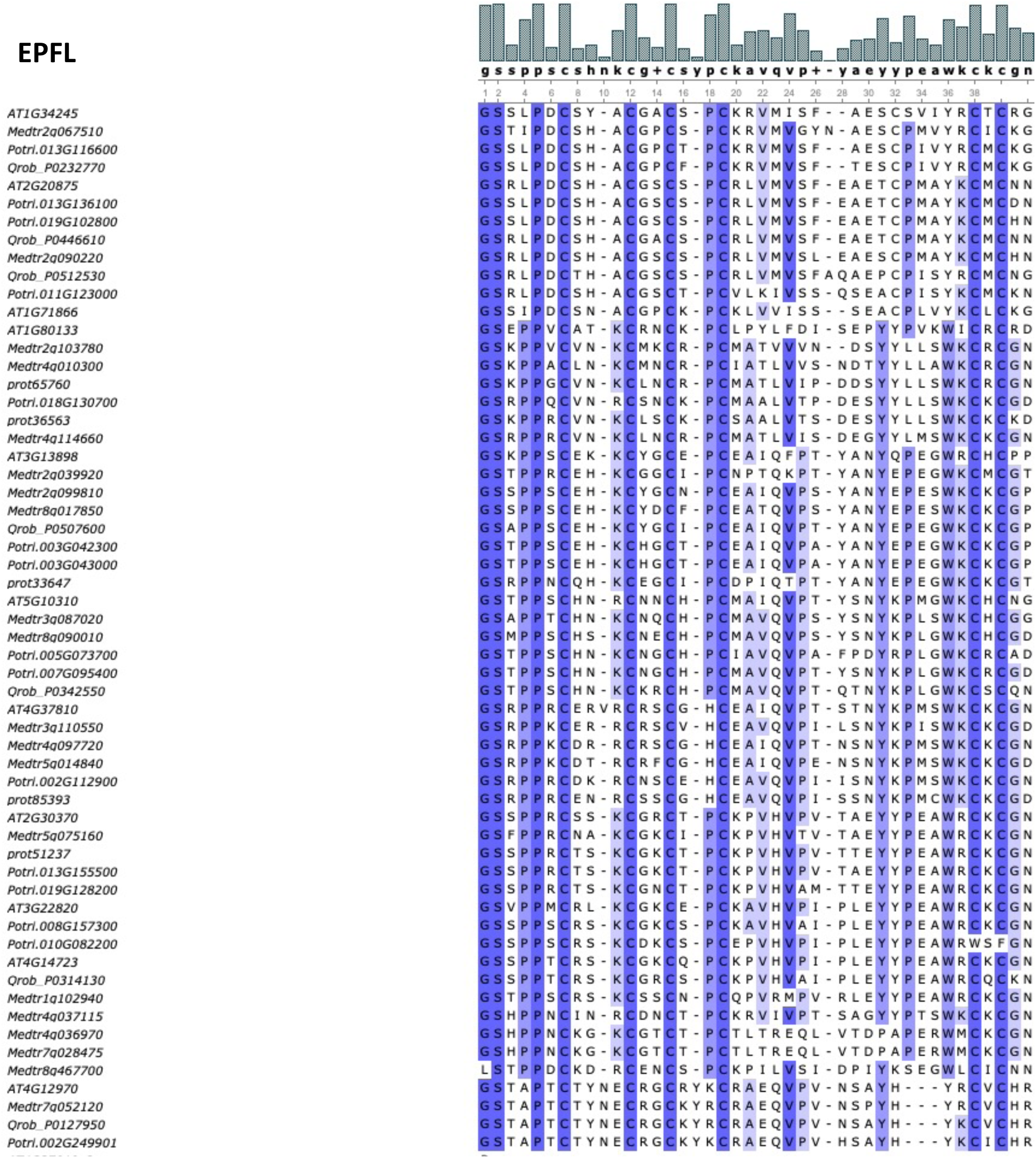
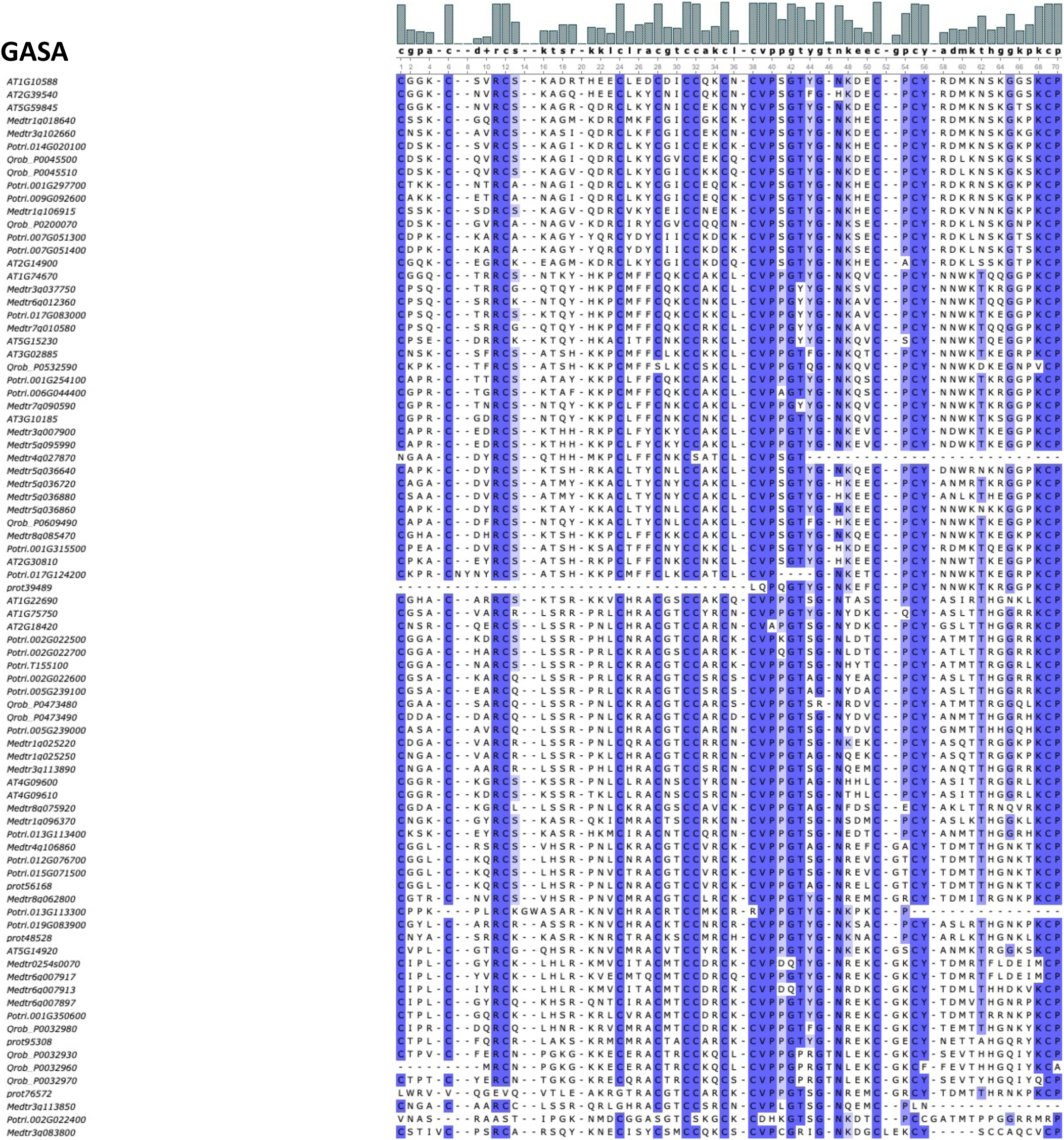
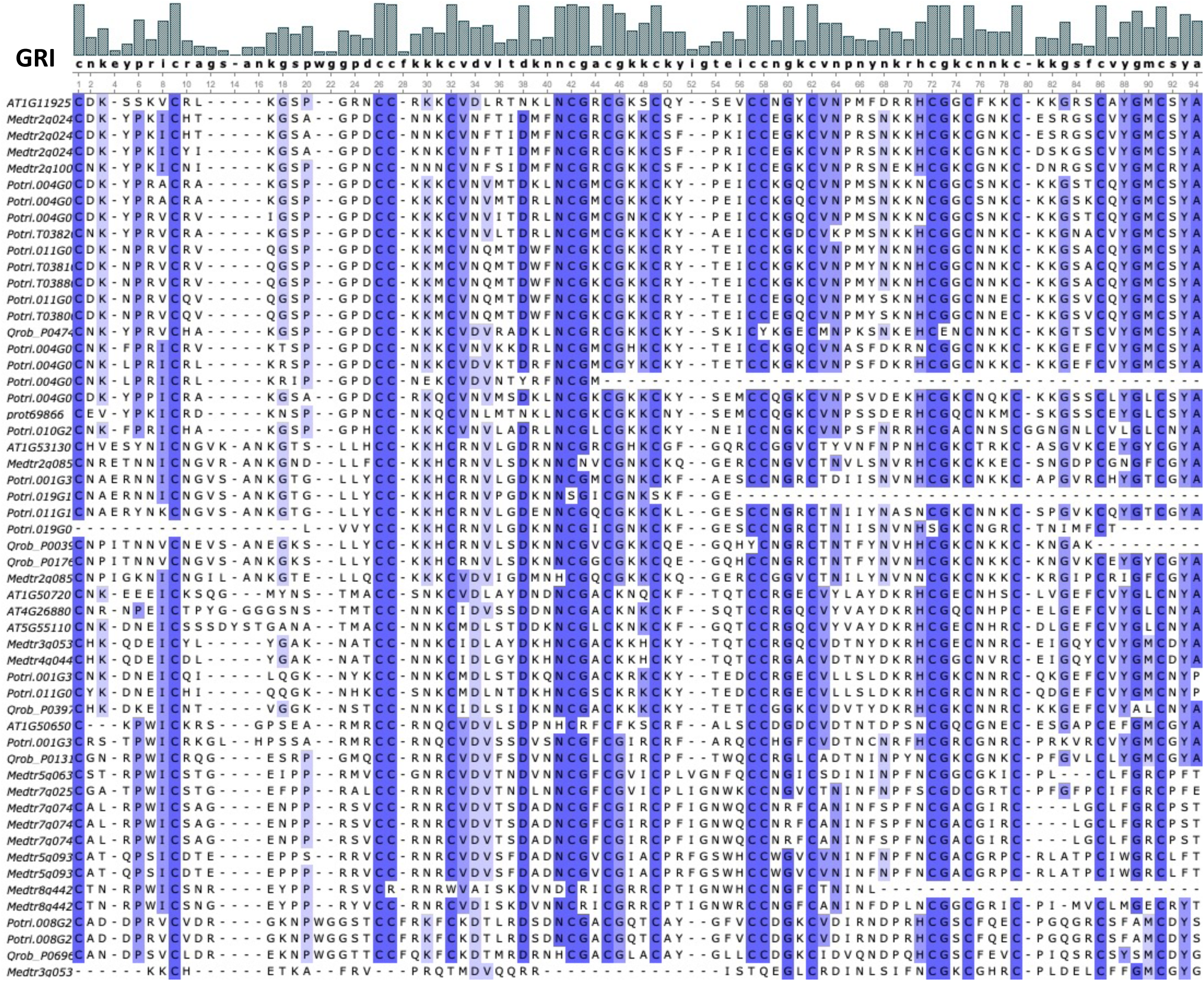
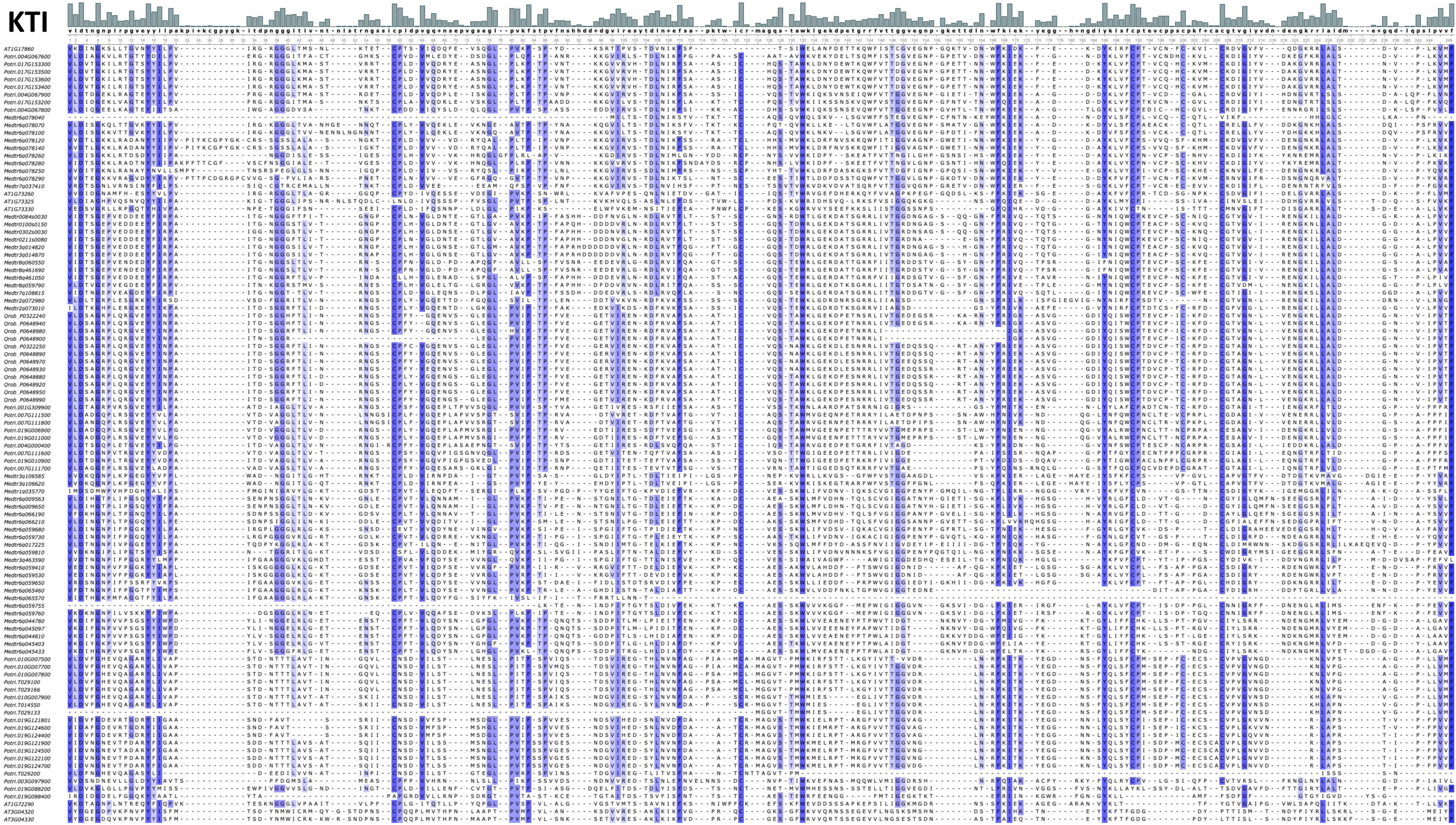
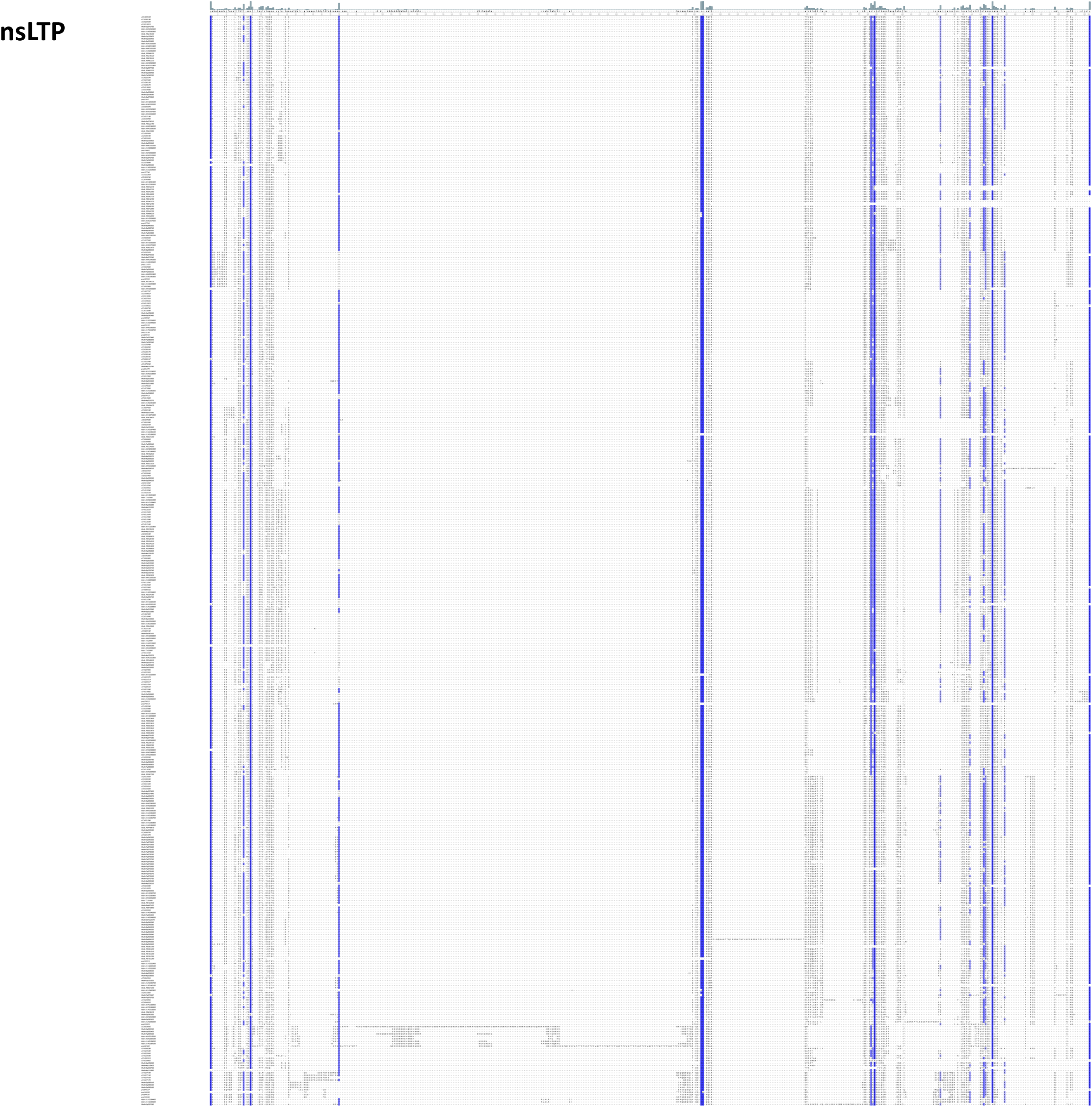
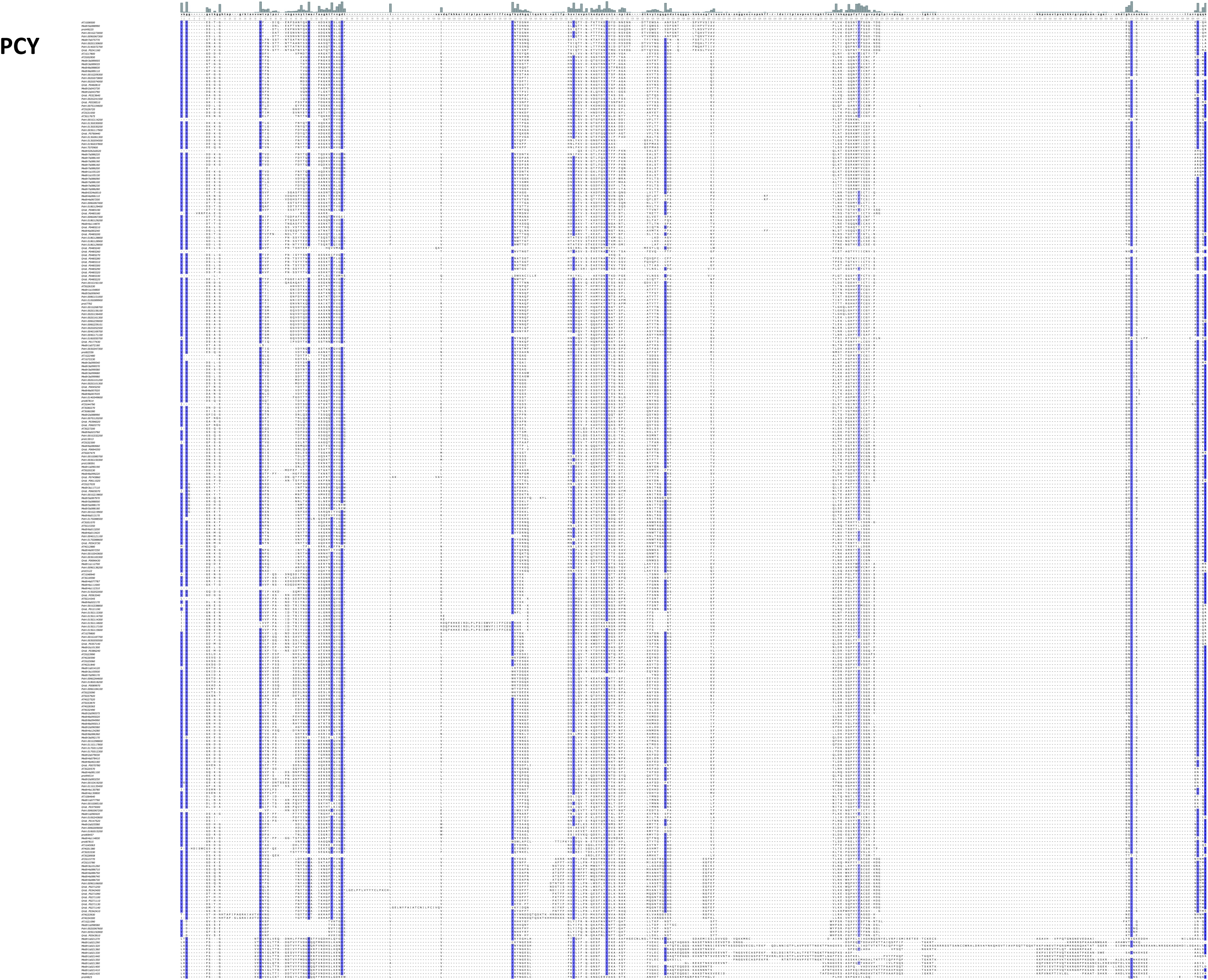
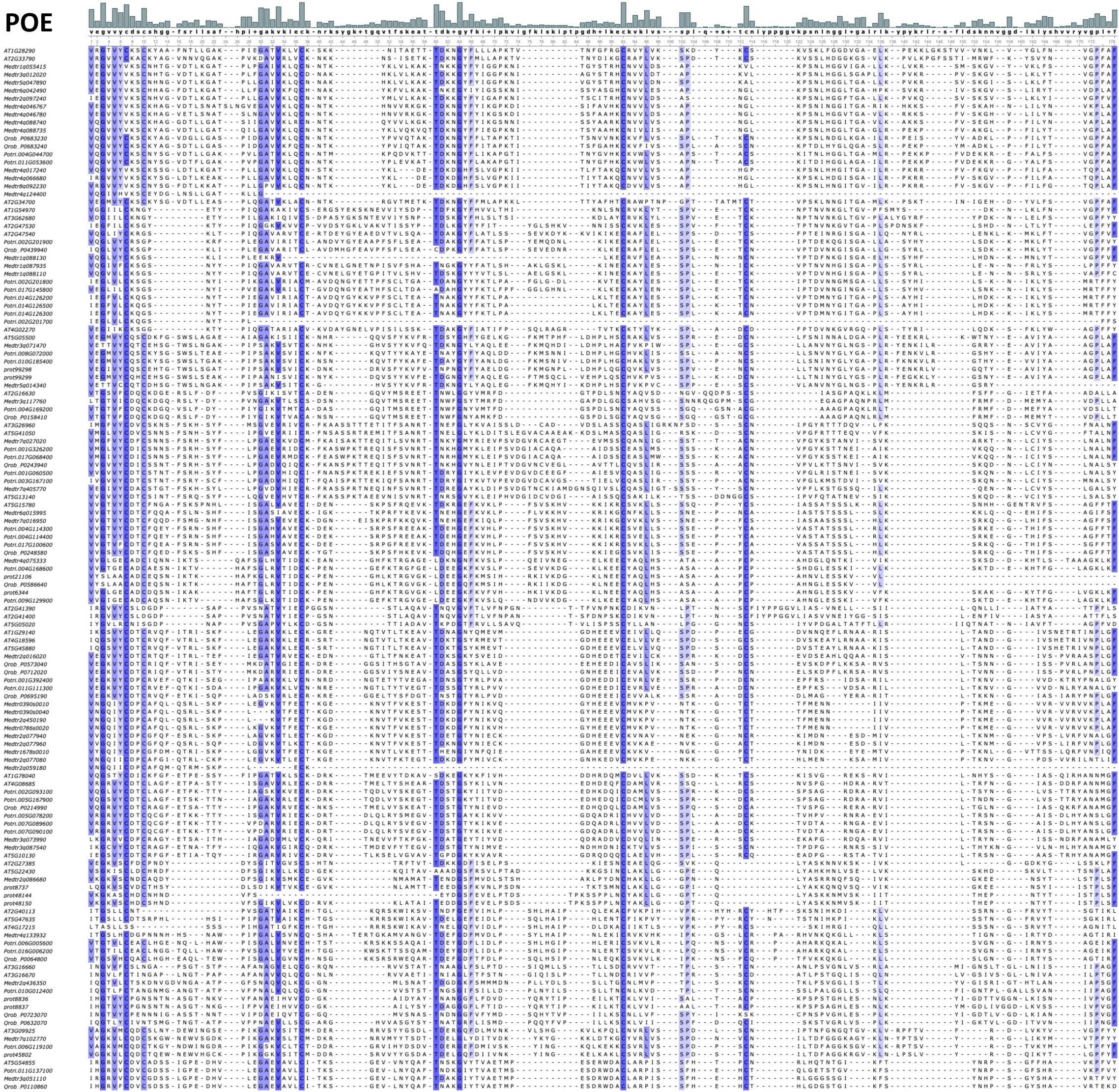
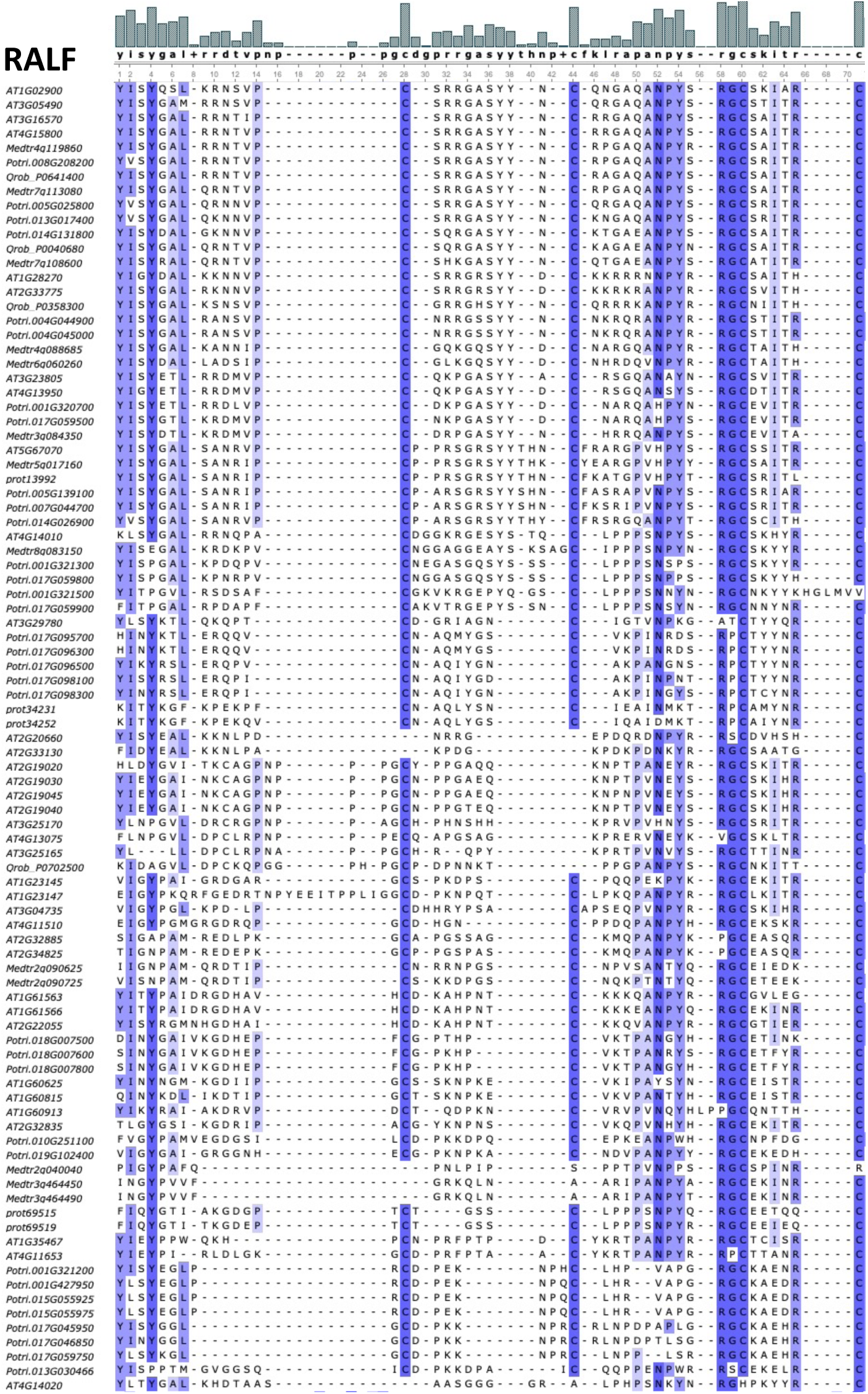
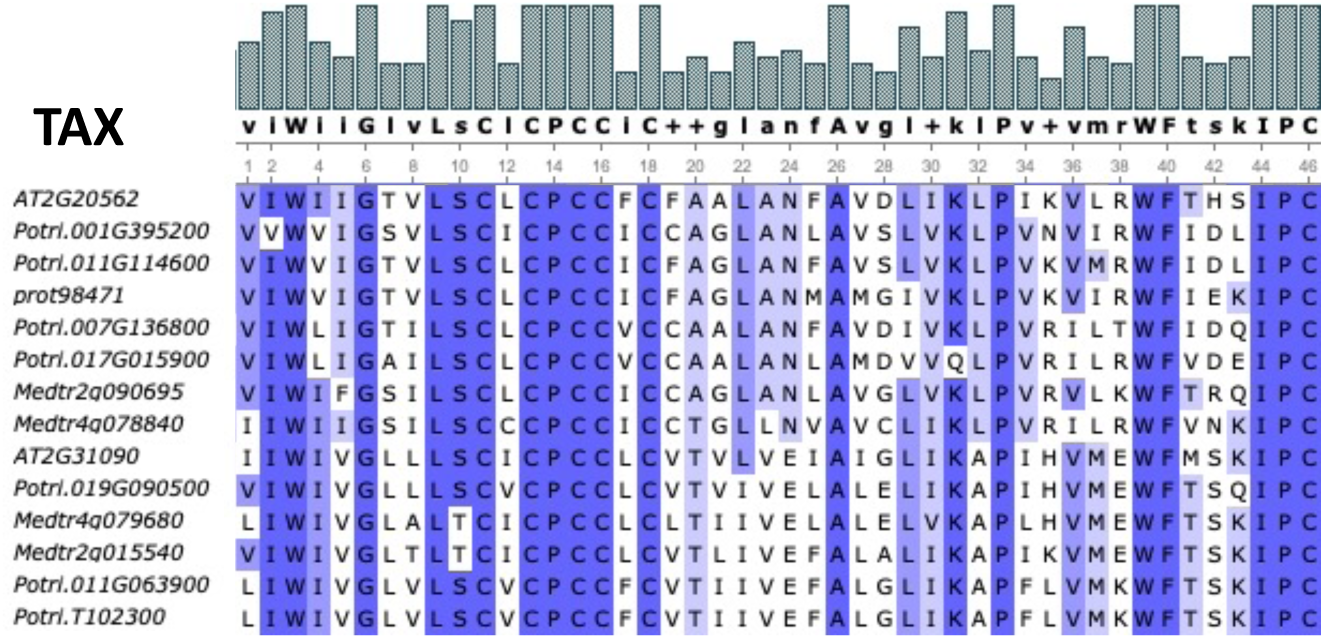
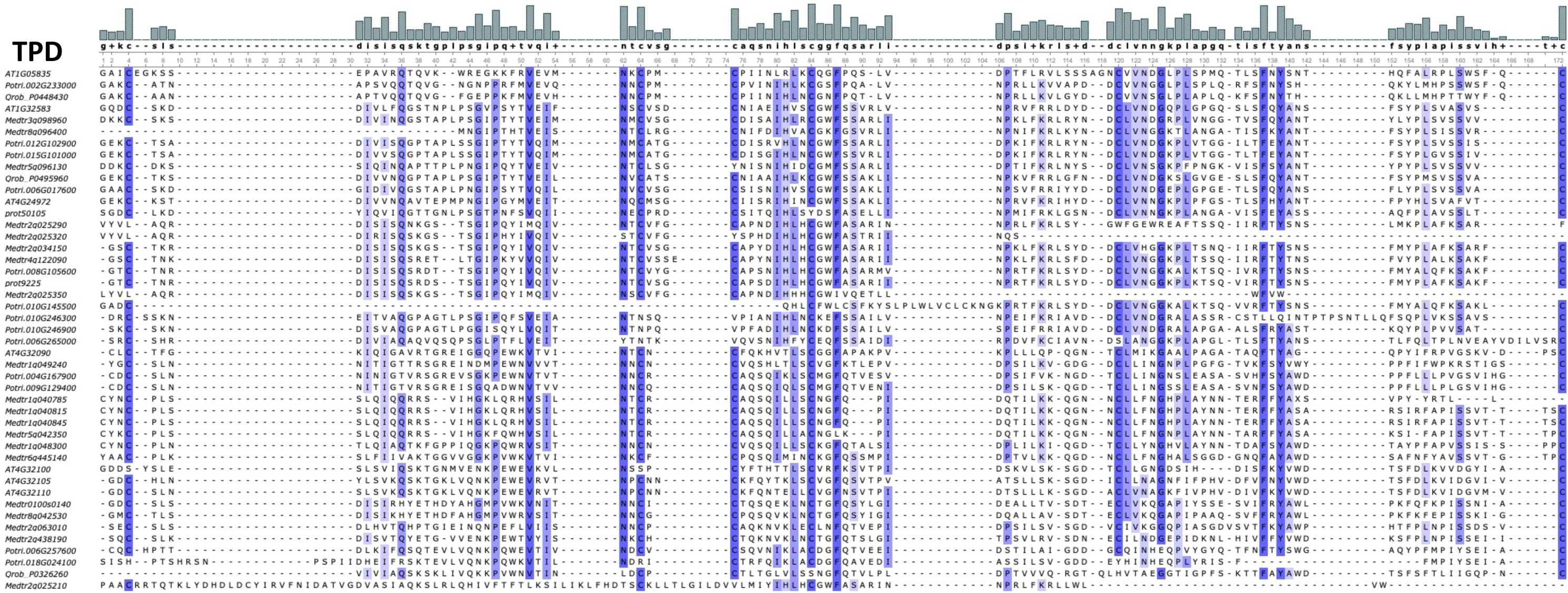
Amino acid alignment of the conserved cysteine patterns of all identified members of the 11 C-rich SSP families. The names of the sequences are the genome identifiers for sequences found from the predicted proteomes, and the identifiers given by the pipeline (Genome Scan ID) for sequences found by *de novo* gene prediction (as in Sup. Table S2). Alignments were produced by Mafft, as described in the Materials and Methods, and visualised using UGENE (Okonechnikov et al., 2012). Blue shading indicates residues conserved in a minimum of 50% of the sequences.

**Sup. Fig. S6.**
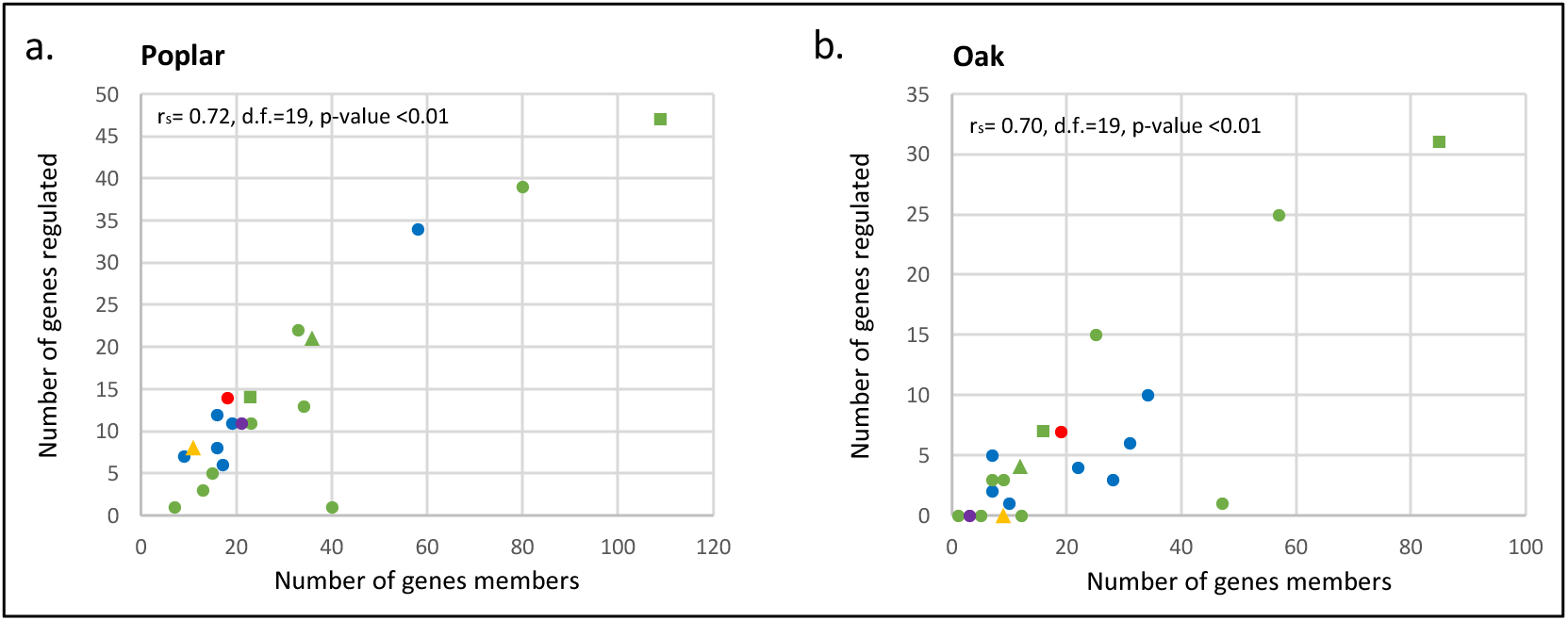
Verification of existing biases associated with SSP family types and functions in SSP regulation by ectomycorrhizal interactions. Regulatory Correlation Plots displaying the number of sequences regulated by ectomycorrhizal interactions in poplar (a) and oak (b) as a function of the number of genes in each of the 21 SSP families. Each point represents one family of SSP. Different colours represent different family types: green for cysteine-rich families (C-rich), blue for Post-Translational Modified peptide families (PTMs), red for the families of peptides produced from functional precursors (FP), purple for the families of peptides produced from small ORFs (sORFs), and yellow for those not entering into any of the previous categories (Other). Different marker shapes represent different family types: points for signalling-peptides, triangles for protease inhibitors, and squares for antimicrobial peptides. The non-parametric Spearman rank-order correlation coefficient (r_S_) was used to verify the statistical significance of correlations between the two variables.

**Sup. Fig. S7.**
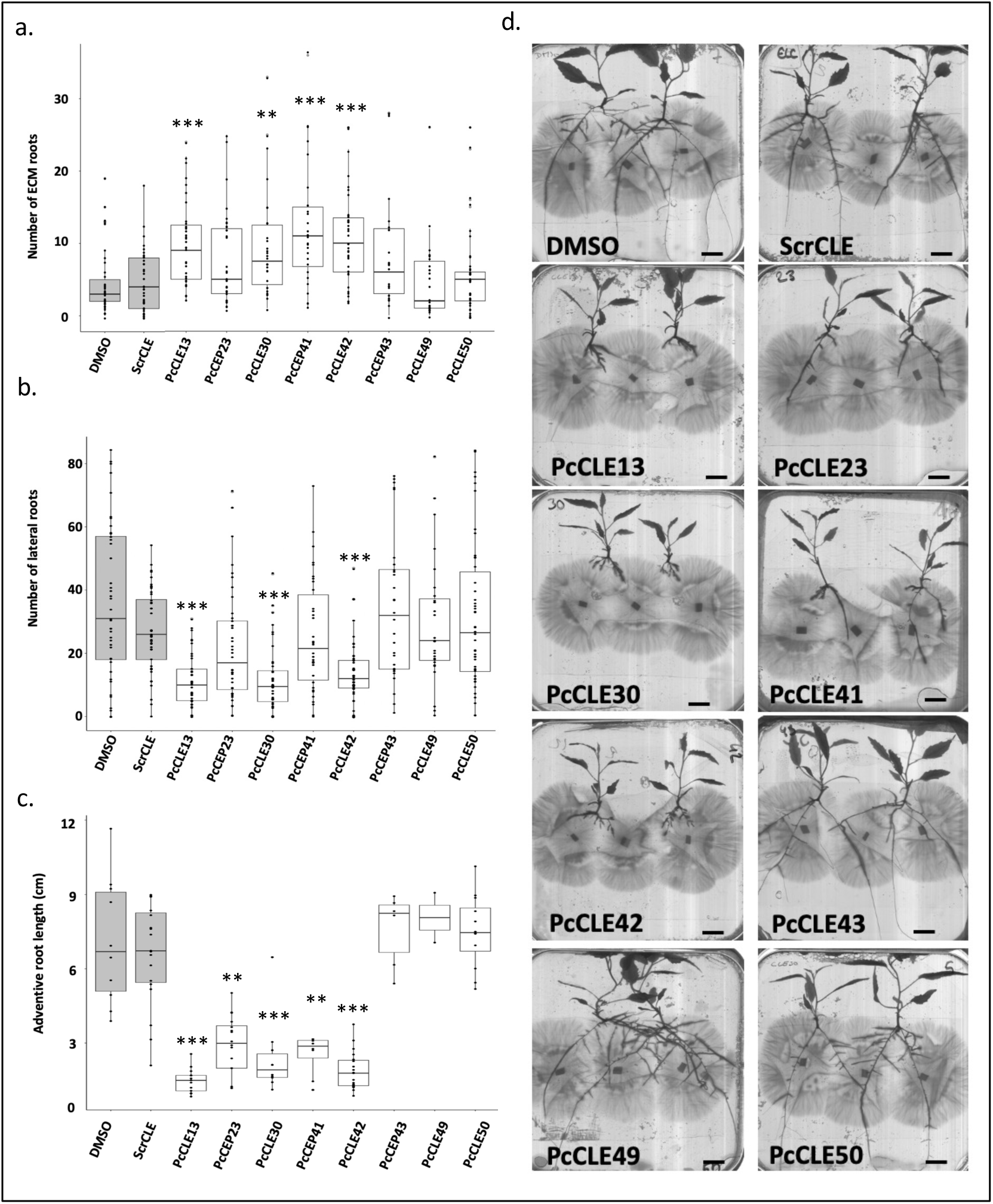
PcCLEs that stimulate ectomycorrhizal associations repress adventive roots growth in *P.* × canescens. a. The number of ectomycorrhizae found between *P. × canescens* and *L. bicolor* co-cultivated on growth media supplemented with 1 µM of PcCLE peptides (PcCLE13, 23, 30, 41, 42, 43, 49, or 50) or with 1 µM of scrambled ScrCLE or with DMSO (solvent) as negative controls. b. The number of lateral roots found in *P. × canescens* co-cultivated with *L. bicolor* on growth media supplemented with 1 µM of PcCLE peptides (PcCLE13, 23, 30, 41, 42, 43, 49, or 50) or with 1 µM of scrambled ScrCLE or with DMSO (solvent) as negative controls. c. Length of adventive roots of *P. × canescens* co-cultivated with *L. bicolor* on growth media supplemented with 1 µM of PcCLE peptides (PcCLE13, 23, 30, 41, 42, 43, 49, or 50) or with 1 µM scrambled ScrCLE or DMSO (solvent) as negative controls. a. b. and d. Asterisks mark treatments statistically significant (* P-value ≤ 0.05; ** P-value ≤ 0.01; *** P-value ≤ 0.001) compared to their controls (DMSO and ScrCLE treatments), based on Kruskal-Wallis one-way analysis of variance and least significant difference (LSD) *post hoc* test corrected with the Bonferroni method, *n* = 25–53. d. Pictures of Petri dishes containing *P. × canescens* co-cultivated with *L. bicolor* on growth media supplemented with 1 µM of PcCLE peptides (PcCLE13, 23, 30, 41, 42, 43, 49, or 50) or with 1 µM of scrambled ScrCLE or with DMSO (solvent) as negative controls. Scale bars represent 1 cm.

**Sup. Tab. S1. Characteristics of the 21 selected SSP families, number of genes member known from the literature and identified by our pipeline in *A. thaliana*, *M. truncatula*, *P. trichocarpa* and *Q. robur*.** For each family, the following information was provided: full name, class, number of conserved cysteines (when applicable), sequence(s) and size(s) of the conserved domain(s) (when applicable), peptide size range, presence of a proteolytic cleavage site, main post-translational modification(s) (Main PTMs), secretion, mode(s) of action, class(es) of receptor(s) (when applicable), example of receptors (when known), and known function(s) and regulation(s) relative to plant nutrition and symbiosis. SSP families are classified into five classes: post-translationally modified (PTM), cysteine-rich (C-rich), functional precursor (FP), short open reading frame (sORF), and those not entering into any of the previous categories (Other). Amino-acids of the SSPs conserved domains are written in upper case, lower case and or as ‘x’ to indicate their conservation at 80%, 60% or < 60%, respectively. For each family, the following numbers are given: number of gene members known from the literature (Known), number of known gene members recovered by the pipeline (Known identified), number of known gene members not predicted by our pipeline (Known not identified) and the number of new gene members identified from our pipeline (New identified) in *A. thaliana*, *M. truncatula*, *P. trichocarpa* and *Q. robur*.

**Sup. Tab. S2. List of SSP-encoding genes identified by the pipeline**. SSP-encoding genes of the 21 families of interest identified from the genomes and predicted proteomes of *A. thaliana*, *M. truncatula*, *P. trichocarpa* and *Q. robur*. For each gene, the following information is given: the SSP family it belongs to, the species it was found in, origin of prediction (predicted proteome or genome), proteome (predicted proteome ID), and genome (Genome ID) identifiers for the sequences found from the predicted proteomes, identifier given by the pipeline (Genome Scan ID) for the sequences found by *de novo* gene prediction, position in genome for the sequences found by *de novo* gene prediction, the prediction method(s) it was predicted by (HMM, MAST and/or BLAST), the number of SSP conserved domains found in the pre-protein (when applicable), whether it was discovered in this study or previously known (Known peptide), its name in the literature when existing (Name), the publication(s) it was found when applicable (Publication(s)), the predicted protein sequence, and other relevant information.

**Sup. Tab. S3. Prediction of localisation.** Prediction of localisation from SignalP V6, DeepLoc V2, and TargetP V2 for all SSP-encoding genes. For each gene the following information is given: the SignalP V6 prediction of signal peptide (signal_peptide), signal peptide start (start pos) and end (end pos) positions and prediction score (score), the DeepLoc V2 prediction of localisation (Localisations) and transit peptide sequence prediction (Signals), localisation prediction scores for each cellular compartment (Cytoplasm, Nucleus, Extracellular, Cell membrane, Mitochondrion, Plastid, Endoplasmic reticulum, Lysosome and Vacuole, Golgi apparatus and Peroxisome), the TargetP V2 prediction of localisation (Prediction) and transit peptides prediction scores (no transit peptide, noTP; Secretory pathway transit peptide, SP; mitochondrial transit peptide, mTP; cTP chloroplast transit peptide, cTP; lumen transit peptide, luTP), and cleavage site position (CS position).

**Sup. Tab. S4. SSP-encoding genes regulated in response to ectomycorrhizal associations in *P. × canescens*.** Were considered regulated SSPs up or down regulated (log2 fold-change ≥ |1|) with a false discovery rate (Padj) corresponding to a Wald test adjusted by the Benjamini–Hochberg procedure for multiple testing inferior to 0.05 in response to the ectomycorrhizal associations of *P. × canescens* with a least one of the four ectomycorrhizal fungi tested *L. bicolor* (Lb), *C. geophylum* (Cg), *P. microcarpus* (Pm) or *A. muscaria* (Am) at one of the two time points (early or late interaction). Roman numbers indicate different genes clusters based on the number of ectomycorrhizal associations inducing or repressing the transcription of the SSP-encoding genes they comprise. Clades I, II, III and IV contained SSP-encoding genes induced respectively by four, three, two or only one ectomycorrhizal interaction. Clades V, VI and VII contained gene oppositely regulated by different ectomycorrhizal interactions, the genes up-regulated by more ectomycorrhizal association in clade V, the genes up- and down regulated by an equal number of ectomycorrhizal associations in clade VI and the genes down-regulated by more ectomycorrhizal associations in clade VII. Clades VIII, IX, X and XI contained SSP-encoding genes repressed respectively by one, two, three or four ectomycorrhizal associations.

**Sup. Tab. S5. SSP-encoding genes regulated in response to ectomycorrhizal associations in *Q. robur*.** Were considered regulated SSPs up or down regulated (log2 fold-change ≥ |1|) with a false discovery rate (Padj) corresponding to a Wald test adjusted by the Benjamini–Hochberg procedure for multiple testing inferior to 0.05 in response to the ectomycorrhizal associations of *Q. robur* with a least one of the three Tuber sp. tested *T. melanosporum* (Tme), *T. aestivum* (Tae) and *T. magnatum* (Tma). Roman numbers indicate different genes clusters based on the number of ectomycorrhizal associations inducing or repressing the transcription of the SSP-encoding genes they comprise. Clusters II, III, and IV contained SSP-encoding genes induced by three, two, or only one ectomycorrhizal interaction, respectively. Clusters V and VI contained genes oppositely regulated by different ectomycorrhizal interactions, genes upregulated by more ectomycorrhizal associations in cluster V, and genes upregulated and downregulated by an equal number of ectomycorrhizal associations in cluster VI. Clusters VIII, IX, and X contained SSP-encoding genes that were repressed by one, two, or three ectomycorrhizal associations.

**Sup. Tab. S6. Database origin and accession numbers of all RNA-Seq datasets from poplar and oak used in this study.** List of accession numbers from the Short Read Archive (SRA) (https://www.ncbi.nlm.nih.gov/sra), GEO (https://www.ncbi.nlm.nih.gov/geo/) and ENA (https://www.ebi.ac.uk/ena/browser/home) databases for all RNA-seq datasets used in this study.

**Sup. Tab. S7. List of primers and melting temperatures used to clone the eight PcCLE-encoding genes selected in this study.** For each PcCLE-encoding gene cloned, the Gene identifier in *P. trichocharpa* (Gene ID) and the two gene identifiers for the tremula (HAP1 ID) and alba (HAP2 ID) haplomes of *P. × canescens*, the forward and reverse primers used, the amplicon size and the melting temperature are given. The version (tremula or alba) cloned from *P. × canescens* is marked in bold red for each PcCLE-encoding gene.

**Sup. Dataset S1**. **Amino acid alignments of all identified members of the 21 SSP families.** The names of the sequences are their genome identifiers when identified from the predicted proteomes, or the identifiers given by the pipeline (Genome Scan ID) when identified by *de novo* gene prediction (as in Sup. Table S2). Alignments were produced by Mafft, as described in the Materials and Methods section.

**Sup. Dataset S2.** Transcriptional regulation of the genes of the 21 symbiosis-related SSP families in response to ectomycorrhizal associations in *P. × canescens*. Relative expression of log2 fold-change of the ratio of gene expression in ectomycorrhiza compared to uncolonized roots in response to association of *P. × canescens* with the ectomycorrhizal fungi *L. bicolor* (Lacbic), *C. geophylum* (Cenge), *P. microcarpus* (Pismi), or *A. muscaria* (Amamu). Early and late are are respectively an early and a late time point of sample collection after contact between the ectomycorrhizal fungi and the host tree (see Material and Method for details). Padj (FDR, false discovery rate), corresponding to a Wald test adjusted by the Benjamini–Hochberg procedure for multiple testing, is given for each gene and condition. Normalised unique read counts from DESEQ2 are provided for all conditions.

**Sup. Dataset S3.** Transcriptional regulation of gene members of the 21 symbiosis-related SSP families in response to ectomycorrhizal associations in *Q. robur*. Relative expression as log2 fold-change of the ratio of genes expression in ectomycorrhiza compared to uncolonized roots in response to association of *Q. robur* with the ectomycorrhizal fungi *T. melanosporum* (Tme), *T. aestivum* (Tae) and *T. magnatum* (Tma). Padj (FDR, false discovery rate), corresponding to a Wald test adjusted by the Benjamini–Hochberg procedure for multiple testing, is also given for each gene and condition. Normalised unique reads count from DESEQ2 are provided for all conditions.

## References

Araya, T., Miyamoto, M., Wibowo, J., Suzuki, A., Kojima, S., Tsuchiya, Y. N., Sawa, S., Fukuda, H., Von Wirén, N., and Takahashi, H. (2014). CLE-CLAVATA1 peptide-receptor signaling module regulates the expansion of plant root systems in a nitrogen-dependent manner. Proceedings of the National Academy of Sciences 111:2029–2034.

Armenteros, J. J. A., Salvatore, M., Emanuelsson, O., Winther, O., Von Heijne, G., Elofsson, A., and Nielsen, H. (2019). Detecting sequence signals in targeting peptides using deep learning. Life science alliance 2.

Bailey, T. L., and Elkan, C. (1994). Fitting a mixture model by expectation maximization to discover motifs in bipolymers Advance Access published 1994.

Bailey, T. L., Johnson, J., Grant, C. E., and Noble, W. S. (2015). The MEME suite. Nucleic acids research 43:W39–W49.

Balbinott, N., and Margis, R. (2022). Unraveling the origin of the structural and functional diversity of plant cystatins. Plant Science 321:111342.

Basso, V., Kohler, A., Miyauchi, S., Singan, V., Guinet, F., Šimura, J., Novák, O., Barry, K. W., Amirebrahimi, M., and Block, J. (2020). An ectomycorrhizal fungus alters sensitivity to jasmonate, salicylate, gibberellin, and ethylene in host roots. Plant, cell & environment 43:1047–1068.

Brand, U., Fletcher, J. C., Hobe, M., Meyerowitz, E. M., and Simon, R. (2000). Dependence of stem cell fate in Arabidopsis on a feedback loop regulated by CLV3 activity. Science 289:617–619.

Breen, S., Williams, S. J., Outram, M., Kobe, B., and Solomon, P. S. (2017). Emerging insights into the functions of pathogenesis-related protein 1. Trends in plant science 22:871–879.

Campbell, L., and Turner, S. R. (2017). A Comprehensive Analysis of RALF Proteins in Green Plants Suggests There Are Two Distinct Functional Groups. Front. Plant Sci. 8.

Cao, J., Li, X., Lv, Y., and Ding, L. (2015). Comparative analysis of the phytocyanin gene family in 10 plant species: a focus on Zea mays. Front. Plant Sci. 6.

Chen, S., Lang, P., Chronis, D., Zhang, S., De Jong, W. S., Mitchum, M. G., and Wang, X. (2015). In planta processing and glycosylation of a nematode CLAVATA3/ENDOSPERM SURROUNDING REGION-like effector and its interaction with a host CLAVATA2-like receptor to promote parasitism. Plant Physiology 167:262–272.

Chen, D., Li, D., Li, Z., Song, Y., Li, Q., Wang, L., Zhou, D., Xie, F., and Li, Y. (2023). Legume nodulation and nitrogen fixation require interaction of DnaJ-like protein and lipid transfer protein. Plant Physiology 193:2164–2179.

Combier, J.-P., Küster, H., Journet, E.-P., Hohnjec, N., Gamas, P., and Niebel, A. (2008). Evidence for the Involvement in Nodulation of the Two Small Putative Regulatory Peptide-Encoding Genes *MtRALFL1* and *MtDVL1*. MPMI 21:1118–1127.

Cope, K. R., Bascaules, A., Irving, T. B., Venkateshwaran, M., Maeda, J., Garcia, K., Rush, T. A., Ma, C., Labbé, J., and Jawdy, S. (2019). The ectomycorrhizal fungus Laccaria bicolor produces lipochitooligosaccharides and uses the common symbiosis pathway to colonize Populus roots. The Plant Cell 31:2386–2410.

Crooks, G. E., Hon, G., Chandonia, J.-M., and Brenner, S. E. (2004). WebLogo: a sequence logo generator. Genome research 14:1188–1190.

Daguerre, Y., Basso, V., Hartmann-Wittulski, S., Schellenberger, R., Meyer, L., Bailly, J., Kohler, A., Plett, J. M., Martin, F., and Veneault-Fourrey, C. (2020). The mutualism effector MiSSP7 of Laccaria bicolor alters the interactions between the poplar JAZ6 protein and its associated proteins. Scientific reports 10:1–16.

de Bang, T. C., Lundquist, P. K., Dai, X., Boschiero, C., Zhuang, Z., Pant, P., Torres-Jerez, I., Roy, S., Nogales, J., Veerappan, V., et al. (2017). Genome-Wide Identification of Medicago Peptides of *Medicago* Peptides Involved in Macronutrient Responses and Nodulation. Plant Physiol. 175:1669–1689.

de Freitas Pereira, M., Veneault-Fourrey, C., Vion, P., Guinet, F., Morin, E., Barry, K. W., Lipzen, A., Singan, V., Pfister, S., and Na, H. (2018). Secretome analysis from the ectomycorrhizal ascomycete Cenococcum geophilum. Frontiers in microbiology 9:141.

Delay, C. (2015). The roles of C-TERMINALLY ENCODED PEPTIDES in Arabidopsis root development Advance Access published 2015.

Delay, C., Imin, N., and Djordjevic, M. A. (2013). CEP genes regulate root and shoot development in response to environmental cues and are specific to seed plants. Journal of Experimental Botany 64:5383–5394.

Deveau, A., Barret, M., Diedhiou, A. G., Leveau, J., de Boer, W., Martin, F., Sarniguet, A., and Frey-Klett, P. (2015). Pairwise transcriptomic analysis of the interactions between the ectomycorrhizal fungus Laccaria bicolor S238N and three beneficial, neutral and antagonistic soil bacteria. Microbial ecology 69:146–159.

Di, Q., Li, Y., Zhang, D., Wu, W., Zhang, L., Zhao, X., Luo, L., and Yu, L. (2022). A novel type of phytosulfokine, PSK-ε, positively regulates root elongation and formation of lateral roots and root nodules in Medicago truncatula. Plant Signaling & Behavior 17:2134672.

Ditengou, F. A., Raudaskoski, M., and Lapeyrie, F. (2003). Hypaphorine, an indole-3-acetic acid antagonist delivered by the ectomycorrhizal fungus Pisolithus tinctorius, induces reorganisation of actin and the microtubule cytoskeleton in Eucalyptus globulus ssp bicostata root hairs. Planta 218:217–225.

Ditengou, F. A., Müller, A., Rosenkranz, M., Felten, J., Lasok, H., Van Doorn, M. M., Legué, V., Palme, K., Schnitzler, J.-P., and Polle, A. (2015). Volatile signalling by sesquiterpenes from ectomycorrhizal fungi reprogrammes root architecture. Nature communications 6:1–9.

Fang, Y., Chang, J., Shi, T., Luo, W., Ou, Y., Wan, D., and Li, J. (2021). Evolution of RGF/GLV/CLEL peptide hormones and their roles in land plant growth and regulation. International Journal of Molecular Sciences 22:13372.

Felten, J., Kohler, A., Morin, E., Bhalerao, R. P., Palme, K., Martin, F., Ditengou, F. A., and Legué, V. (2009). The ectomycorrhizal fungus Laccaria bicolor stimulates lateral root formation in poplar and Arabidopsis through auxin transport and signaling. Plant Physiology 151:1991–2005.

Fiers, M., Golemiec, E., Xu, J., van der Geest, L., Heidstra, R., Stiekema, W., and Liu, C.-M. (2005). The 14–Amino Acid CLV3, CLE19, and CLE40 Peptides Trigger Consumption of the Root Meristem in Arabidopsis through a CLAVATA2-Dependent Pathway. The Plant Cell 17:2542–2553.

Fleury, C., Gracy, J., Gautier, M.-F., Pons, J.-L., Dufayard, J.-F., Labesse, G., Ruiz, M., and De Lamotte, F. (2019). Comprehensive classification of the plant non-specific lipid transfer protein superfamily towards its sequence–structure–function analysis. PeerJ 7:e7504.

Frickey, T., and Lupas, A. (2004). CLANS: a Java application for visualizing protein families based on pairwise similarity. Bioinformatics 20:3702–3704.

Furumizu, C., Krabberød, A. K., Hammerstad, M., Alling, R. M., Wildhagen, M., Sawa, S., and Aalen, R. B. (2021). The sequenced genomes of nonflowering land plants reveal the innovative evolutionary history of peptide signaling. The Plant Cell 33:2915–2934.

Gao, H., Ma, K., Ji, G., Pan, L., and Zhou, Q. (2022). Lipid transfer proteins involved in plant–pathogen interactions and their molecular mechanisms. Molecular Plant Pathology 23:1815–1829.

Garcia, K., Delaux, P.-M., Cope, K. R., and Ané, J.-M. (2015). Molecular signals required for the establishment and maintenance of ectomycorrhizal symbioses. New Phytologist 208:79–87.

Gasteiger, E., Gattiker, A., Hoogland, C., Ivanyi, I., Appel, R. D., and Bairoch, A. (2003). ExPASy: the proteomics server for in-depth protein knowledge and analysis. Nucleic acids research 31:3784–3788.

Gautrat, P., Laffont, C., and Frugier, F. (2020). Compact root architecture 2 promotes root competence for nodulation through the miR2111 systemic effector. Current Biology 30:1339–1345. e3.

Gholami, A. (2013). Identification of Potential Regulators of Jasmonate-Modulated Secondary Metabolism in Medicago truncatula Advance Access published 2013.

Ghorbani, S., Lin, Y.-C., Parizot, B., Fernandez, A., Njo, M. F., Van de Peer, Y., Beeckman, T., and Hilson, P. (2015). Expanding the repertoire of secretory peptides controlling root development with comparative genome analysis and functional assays. Journal of experimental botany 66:5257–5269.

Goad, D. M., Zhu, C., and Kellogg, E. A. (2017). Comprehensive identification and clustering of CLV3/ESR-related (CLE) genes in plants finds groups with potentially shared function. New Phytologist 216:605–616.

Grigoriev, I. V., Nikitin, R., Haridas, S., Kuo, A., Ohm, R., Otillar, R., Riley, R., Salamov, A., Zhao, X., and Korzeniewski, F. (2014). MycoCosm portal: gearing up for 1000 fungal genomes. Nucleic acids research 42:D699–D704.

Guindon, S., Dufayard, J.-F., Lefort, V., Anisimova, M., Hordijk, W., and Gascuel, O. (2010). New algorithms and methods to estimate maximum-likelihood phylogenies: assessing the performance of PhyML 3.0. Systematic biology 59:307–321.

Guo, P., Yoshimura, A., Ishikawa, N., Yamaguchi, T., Guo, Y., and Tsukaya, H. (2015). Comparative analysis of the RTFL peptide family on the control of plant organogenesis. J Plant Res 128:497–510.

Guo, X., Wang, J., Gardner, M., Fukuda, H., Kondo, Y., Etchells, J. P., Wang, X., and Mitchum, M. G. (2017). Identification of cyst nematode B-type CLE peptides and modulation of the vascular stem cell pathway for feeding cell formation. PLoS Pathogens 13:e1006142.

Guo, C., Wang, Q., Li, Z., Sun, J., Zhang, Z., Li, X., and Guo, Y. (2021). Bioinformatics and expression analysis of IDA-Like genes reveal their potential functions in flower abscission and stress response in tobacco (Nicotiana tabacum L.). Frontiers in Genetics 12:670794.

Gutiérrez-Alanís, D., Yong-Villalobos, L., Jiménez-Sandoval, P., Alatorre-Cobos, F., Oropeza-Aburto, A., Mora-Macías, J., Sánchez-Rodríguez, F., Cruz-Ramírez, A., and Herrera-Estrella, L. (2017). Phosphate Starvation-Dependent Iron Mobilization Induces CLE14 Expression to Trigger Root Meristem Differentiation through CLV2/PEPR2 Signaling. Developmental Cell 41:555–570.e3.

Han, Z., Xiong, D., Schneiter, R., and Tian, C. (2023). The function of plant PR1 and other members of the CAP protein superfamily in plant–pathogen interactions. Molecular plant pathology 24:651–668.

Hanada, K., Zhang, X., Borevitz, J. O., Li, W.-H., and Shiu, S.-H. (2007). A large number of novel coding small open reading frames in the intergenic regions of the Arabidopsis thaliana genome are transcribed and/or under purifying selection. Genome research 17:632–640.

Hastwell, A. H., Gresshoff, P. M., and Ferguson, B. J. (2014). The structure and activity of nodulation-suppressing CLE peptide hormones of legumes. Functional Plant Biology 42:229–238.

Hastwell, A. H., de Bang, T. C., Gresshoff, P. M., and Ferguson, B. J. (2017). CLE peptide-encoding gene families in Medicago truncatula and Lotus japonicus, compared with those of soybean, common bean and Arabidopsis. Scientific Reports 7:9384.

Hastwell, A. H., Corcilius, L., Williams, J. T., Gresshoff, P. M., Payne, R. J., and Ferguson, B. J. (2019). Triarabinosylation is required for nodulation-suppressive CLE peptides to systemically inhibit nodulation in Pisum sativum. Plant, Cell & Environment 42:188–197.

He, L., Chen, X., Xu, M., Liu, T., Zhang, T., Li, J., Yang, J., Chen, J., and Zhong, K. (2021). Genome-wide identification and characterization of the cystatin gene family in bread wheat (Triticum aestivum L.). International Journal of Molecular Sciences 22:10264.

Hirakawa, Y., Kondo, Y., and Fukuda, H. (2010). TDIF peptide signaling regulates vascular stem cell proliferation via the WOX4 homeobox gene in Arabidopsis. The Plant Cell 22:2618–2629.

Hortal, S., Plett, K. L., Plett, J. M., Cresswell, T., Johansen, M., Pendall, E., and Anderson, I. C. (2017). Role of plant–fungal nutrient trading and host control in determining the competitive success of ectomycorrhizal fungi. The ISME journal 11:2666.

Hou, S., Wang, X., Chen, D., Yang, X., Wang, M., Turrà, D., Di Pietro, A., and Zhang, W. (2014). The secreted peptide PIP1 amplifies immunity through receptor-like kinase 7. PLoS pathogens 10:e1004331.

Hu, X.-L., Lu, H., Hassan, M. M., Zhang, J., Yuan, G., Abraham, P. E., Shrestha, H. K., Villalobos Solis, M. I., Chen, J.-G., and Tschaplinski, T. J. (2021). Advances and perspectives in discovery and functional analysis of small secreted proteins in plants. Horticulture research 8.

Hu, X.-L., Zhang, J., Kaundal, R., Kataria, R., Labbé, J. L., Mitchell, J. C., Tschaplinski, T. J., Tuskan, G. A., Cheng, Z.-M., and Yang, X. (2022). Diversity and conservation of plant small secreted proteins associated with arbuscular mycorrhizal symbiosis. Horticulture research 9:uhac043.

Ikeuchi, M., Yamaguchi, T., Kazama, T., Ito, T., Horiguchi, G., and Tsukaya, H. (2011). ROTUNDIFOLIA4 regulates cell proliferation along the body axis in Arabidopsis shoot. Plant and Cell Physiology 52:59–69.

Imin, N., Mohd-Radzman, N. A., Ogilvie, H. A., and Djordjevic, M. A. (2013). The peptide-encoding CEP1 gene modulates lateral root and nodule numbers in Medicago truncatula. Journal of experimental botany 64:5395–5409.

Imin, N., Patel, N., Corcilius, L., Payne, R. J., and Djordjevic, M. A. (2018). CLE peptide tri-arabinosylation and peptide domain sequence composition are essential for SUNN-dependent autoregulation of nodulation in Medicago truncatula. New Phytologist 218:73–80.

Jiang, W., Chen, S.-Y., Wang, H., Li, D.-Z., and Wiens, J. J. (2014). Should genes with missing data be excluded from phylogenetic analyses? Molecular phylogenetics and evolution 80:308–318.

Jiménez-López, J. C., Rodríguez-García, M. I., and de Dios Alché, J. (2011). Systematic and Phylogenetic Analysis of the Ole e 1 Pollen Protein Family Members in Plants. In Systems and Computational Biology - Bioinformatics and Computational Modeling (ed. Yang, N.-S.), p. InTech.

Kang, H., Chen, X., Kemppainen, M., Pardo, A. G., Veneault-Fourrey, C., Kohler, A., and Martin, F. M. (2020). The small secreted effector protein MiSSP7. 6 of Laccaria bicolor is required for the establishment of ectomycorrhizal symbiosis. Environmental Microbiology 22:1435–1446.

Karlo, M., Boschiero, C., Landerslev, K. G., Blanco, G. S., Wen, J., Mysore, K. S., Dai, X., Zhao, P. X., and de Bang, T. C. (2020). The CLE53–SUNN genetic pathway negatively regulates arbuscular mycorrhiza root colonization in Medicago truncatula. Journal of experimental botany 71:4972–4984.

Katoh, K., and Standley, D. M. (2013). MAFFT multiple sequence alignment software version 7: improvements in performance and usability. Molecular biology and evolution 30:772–780.

Keerthana, K., Ramakrishnan, M., Ahmad, Z., Amali, P., Vijayakanth, V., and Wei, Q. (2025). Root-derived small peptides: Key regulators of plant development, stress resilience, and nutrient acquisition. Plant Science Advance Access published 2025.

Keller, O., Kollmar, M., Stanke, M., and Waack, S. (2011). A novel hybrid gene prediction method employing protein multiple sequence alignments. Bioinformatics 27:757–763.

Kim, M.-J., Jeon, B. W., Oh, E., Seo, P. J., and Kim, J. (2021). Peptide signaling during plant reproduction. Trends in Plant Science 26:822–835.

Kohler, A., Kuo, A., Nagy, L. G., Morin, E., Barry, K. W., Buscot, F., Canbäck, B., Choi, C., Cichocki, N., Clum, A., et al. (2015). Convergent losses of decay mechanisms and rapid turnover of symbiosis genes in mycorrhizal mutualists. Nat Genet 47.

Kucukoglu, M., Chaabouni, S., Zheng, B., Mähönen, A. P., Helariutta, Y., and Nilsson, O. (2020). Peptide encoding Populus CLV3/ESR-RELATED 47 (PttCLE47) promotes cambial development and secondary xylem formation in hybrid aspen. New Phytologist 226:75–85.

Labbé, J., Jorge, V., Kohler, A., Vion, P., Marçais, B., Bastien, C., Tuskan, G. A., Martin, F., and Le Tacon, F. (2011). Identification of quantitative trait loci affecting ectomycorrhizal symbiosis in an interspecific F1 poplar cross and differential expression of genes in ectomycorrhizas of the two parents: Populus deltoides and Populus trichocarpa. Tree Genetics & Genomes 7:617–627.

Labbé, J., Muchero, W., Czarnecki, O., Wang, J., Wang, X., Bryan, A. C., Zheng, K., Yang, Y., Xie, M., and Zhang, J. (2019). Mediation of plant–mycorrhizal interaction by a lectin receptor-like kinase. Nature Plants 5:676–680.

Laffont, C., Ivanovici, A., Gautrat, P., Brault, M., Djordjevic, M. A., and Frugier, F. (2020). The NIN transcription factor coordinates CEP and CLE signaling peptides that regulate nodulation antagonistically. Nature communications 11:1–13.

Lamesch, P., Berardini, T. Z., Li, D., Swarbreck, D., Wilks, C., Sasidharan, R., Muller, R., Dreher, K., Alexander, D. L., and Garcia-Hernandez, M. (2012). The Arabidopsis Information Resource (TAIR): improved gene annotation and new tools. Nucleic acids research 40:D1202–D1210.

Le Marquer, M., Bécard, G., and Frei dit Frey, N. (2019). Arbuscular mycorrhizal fungi possess a CLAVATA3/embryo surrounding region-related gene that positively regulates symbiosis. New Phytologist 222:1030–1042.

Lease, K. A., and Walker, J. C. (2006). The Arabidopsis unannotated secreted peptide database, a resource for plant peptidomics. Plant physiology 142:831–838.

Lefort, V., Longueville, J.-E., and Gascuel, O. (2017). SMS: smart model selection in PhyML. Molecular biology and evolution 34:2422–2424.

Li, Y. L., Dai, X. R., Yue, X., Gao, X.-Q., and Zhang, X. S. (2014). Identification of small secreted peptides (SSPs) in maize and expression analysis of partial SSP genes in reproductive tissues. Planta 240:713–728.

Li, Q., Li, M., Zhang, D., Yu, L., Yan, J., and Luo, L. (2020). The peptide-encoding MtRGF3 gene negatively regulates nodulation of Medicago truncatula. Biochemical and Biophysical Research Communications 523:66–71.

Li, Q., Shan, D., Zheng, W., Wang, Y., Lin, Z., Jin, H., Ding, A., Yan, J., Yu, L., and Luo, L. (2023). MtRGF3 peptide activates defense responses and represses the expressions of nodulation signaling genes in Medicago truncatula: activation of defense response by MtRGF3p. Acta Biochimica et Biophysica Sinica 55:1319.

Liu, Y., Yang, S., Song, Y., Men, S., and Wang, J. (2016). Gain-of-function analysis of poplar CLE genes in Arabidopsis by exogenous application and over-expression assays. Journal of experimental botany 67:2309–2324.

Liu, Y., Zhang, F., Devireddy, A. R., Ployet, R. A., Rush, T. A., Lu, H., Hassan, M. M., Yuan, G., Rajput, R., and Islam, M. T. (2024). A small secreted protein serves as a plant-derived effector mediating symbiosis between Populus and Laccaria bicolor. Horticulture Research 11:uhae232.

Love, M., Anders, S., and Huber, W. (2014). Differential analysis of count data–the DESeq2 package. Genome Biol 15:10–1186.

Luo, S., Hu, W., Wang, Y., Liu, B., Yan, H., and Xiang, Y. (2018). Genome-wide identification, classification, and expression of phytocyanins in Populus trichocarpa. Planta 247:1133–1148.

Ma, Y., Zhao, Q., Lu, M.-Z., and Wang, J. (2011). Kunitz-type trypsin inhibitor gene family in Arabidopsis and Populus trichocarpa and its expression response to wounding and herbivore in Populus nigra. Tree Genetics & Genomes 7:431–441.

Marqués-Gálvez, J. E., Veneault-Fourrey, C., and Kohler, A. (2022). Ectomycorrhizal symbiosis: from genomics to trans-kingdom molecular communication and signaling. In Microbial cross-talk in the rhizosphere, pp. 273–296. Springer.

Marqués-Gálvez, J. E., Pandharikar, G., Basso, V., Kohler, A., Lackus, N. D., Barry, K., Keymanesh, K., Johnson, J., Singan, V., Grigoriev, I. V., et al. (2024). Populus MYC2 orchestrates root transcriptional reprogramming of defence pathway to impair Laccaria bicolor ectomycorrhizal development. New Phytologist 242:658–674.

Marqués-Gálvez, J. E., de Freitas Pereira, M., Nehls, U., Ruytinx, J., Barry, K., Peter, M., Martin, F., Grigoriev, I. V., Veneault-Fourrey, C., and Kohler, A. (2025). Comparative transcriptomics uncovers plant and fungal genetic determinants of mycorrhizal compatibility. bioRxiv Advance Access published January 1, 2025, doi:10.1101/2025.04.11.648352.

Martin, F., Kohler, A., Murat, C., Veneault-Fourrey, C., and Hibbett, D. S. (2016). Unearthing the roots of ectomycorrhizal symbioses. Nat Rev Microbiol 14.

Martinez, M., and Diaz, I. (2008). The origin and evolution of plant cystatins and their target cysteine proteinases indicate a complex functional relationship. BMC Evolutionary Biology 8:1–12.

Martínez, M., Cambra, I., González-Melendi, P., Santamaría, M. E., and Díaz, I. (2012). C1A cysteine-proteases and their inhibitors in plants. Physiologia Plantarum 145:85–94.

Massoumou, M., van Tuinen, D., Chatagnier, O., Arnould, C., Brechenmacher, L., Sanchez, L., Selim, S., Gianinazzi, S., and Gianinazzi-Pearson, V. (2007). Medicago truncatula gene responses specific to arbuscular mycorrhiza interactions with different species and genera of Glomeromycota. Mycorrhiza 17:223–234.

Matsubayashi, Y. (2014). Posttranslationally modified small-peptide signals in plants. Annual review of plant biology 65:385–413.

Mens, C., Hastwell, A. H., Su, H., Gresshoff, P. M., Mathesius, U., and Ferguson, B. J. (2021). Characterisation of Medicago truncatula CLE34 and CLE35 in nitrate and rhizobia regulation of nodulation. New Phytologist 229:2525–2534.

Miyauchi, S., Kiss, E., Kuo, A., Drula, E., Kohler, A., Sánchez-García, M., Morin, E., Andreopoulos, B., Barry, K. W., and Bonito, G. (2020). Large-scale genome sequencing of mycorrhizal fungi provides insights into the early evolution of symbiotic traits. Nature communications 11:5125.

Mohd-Radzman, N. A., Laffont, C., Ivanovici, A., Patel, N., Reid, D., Stougaard, J., Frugier, F., Imin, N., and Djordjevic, M. A. (2016). Different pathways act downstream of the CEP peptide receptor CRA2 to regulate lateral root and nodule development. Plant Physiology 171:2536–2548.

Mortier, V., Den Herder, G., Whitford, R., Van de Velde, W., Rombauts, S., D’haeseleer, K., Holsters, M., and Goormachtig, S. (2010). CLE peptides control Medicago truncatula nodulation locally and systemically. Plant Physiology 153:222–237.

Müller, L. M., Flokova, K., Schnabel, E., Sun, X., Fei, Z., Frugoli, J., Bouwmeester, H. J., and Harrison, M. J. (2019). A CLE–SUNN module regulates strigolactone content and fungal colonization in arbuscular mycorrhiza. Nature Plants 5:933–939.

Murashige, T., and Skoog, F. (1962). A revised medium for rapid growth and bio assays with tobacco tissue cultures. Physiologia plantarum 15:473–497.

Myburg, A. A., Grattapaglia, D., Tuskan, G. A., Hellsten, U., Hayes, R. D., Grimwood, J., Jenkins, J., Lindquist, E., Tice, H., and Bauer, D. (2014). The genome of Eucalyptus grandis. Nature 510:356–362.

Nahirñak, V., Almasia, N. I., Lia, V. V., Hopp, H. E., and Vazquez Rovere, C. (2024). Unveiling the defensive role of Snakin-3, a member of the subfamily III of Snakin/GASA peptides in potatoes. Plant Cell Reports 43:47.

Nakagami, S., Kajiwara, T., Tsuda, K., and Sawa, S. (2024). CLE peptide signaling in plant-microbe interactions. Frontiers in Plant Science 15:1481650.

Nishida, H., Handa, Y., Tanaka, S., Suzaki, T., and Kawaguchi, M. (2016). Expression of the CLE-RS3 gene suppresses root nodulation in Lotus japonicus. Journal of plant research 129:909–919.

Oelkers, K., Goffard, N., Weiller, G. F., Gresshoff, P. M., Mathesius, U., and Frickey, T. (2008). Bioinformatic analysis of the CLE signaling peptide family. BMC Plant Biology 8:1–15.

Ohyama, K., Ogawa, M., and Matsubayashi, Y. (2008). Identification of a biologically active, small, secreted peptide in Arabidopsis by in silico gene screening, followed by LC-MS-based structure analysis. The Plant Journal 55:152–160.

Okamoto, S., Ohnishi, E., Sato, S., Takahashi, H., Nakazono, M., Tabata, S., and Kawaguchi, M. (2008). Nod factor/nitrate-induced CLE genes that drive HAR1-mediated systemic regulation of nodulation. Plant and Cell Physiology 50:67–77.

Okamoto, S., Shinohara, H., Mori, T., Matsubayashi, Y., and Kawaguchi, M. (2013). Root-derived CLE glycopeptides control nodulation by direct binding to HAR1 receptor kinase. Nature communications 4:2191.

Okonechnikov, K., Golosova, O., Fursov, M., and Team, U. (2012). Unipro UGENE: a unified bioinformatics toolkit. Bioinformatics 28:1166–1167.

Oldroyd, G. E. (2013). Speak, friend, and enter: signalling systems that promote beneficial symbiotic associations in plants. Nature Reviews Microbiology 11:252–263.

Olsson, V., Joos, L., Zhu, S., Gevaert, K., Butenko, M. A., and De Smet, I. (2019). Look closely, the beautiful may be small: precursor-derived peptides in plants. Annual review of plant biology 70:153–186.

Onrubia, M., Pollier, J., Vanden Bossche, R., Goethals, M., Gevaert, K., Moyano, E., Vidal-Limon, H., Cusidó, R. M., Palazón, J., and Goossens, A. (2014). Taximin, a conserved plant-specific peptide is involved in the modulation of plant-specialized metabolism. Plant Biotechnol J 12:971–983.

Pan, B., Sheng, J., Sun, W., Zhao, Y., Hao, P., and Li, X. (2012). OrysPSSP: a comparative platform for small secreted proteins from rice and other plants. Nucleic acids research 41:D1192–D1198.

Pedinotti, L., Teyssendier de la Serve, J., Roudaire, T., San Clemente, H., Aguilar, M., Kohlen, W., Frugier, F., and Frei dit Frey, N. (2024). The CEP peptide-CRA2 receptor module promotes arbuscular mycorrhizal symbiosis. Current Biology Advance Access published October 21, 2024, doi:10.1016/j.cub.2024.09.058.

Philippe, R. N., Ralph, S. G., Külheim, C., Jancsik, S. I., and Bohlmann, J. (2009). Poplar defense against insects: genome analysis, full-length cDNA cloning, and transcriptome and protein analysis of the poplar Kunitz-type protease inhibitor family. New Phytologist 184:865–884.

Plett, J. M., Kemppainen, M., Kale, S. D., Kohler, A., Legué, V., Brun, A., Tyler, B. M., Pardo, A. G., and Martin, F. (2011). A secreted effector protein of Laccaria bicolor is required for symbiosis development. Current Biology 21:1197–1203.

Plett, J. M., Daguerre, Y., Wittulsky, S., Vayssières, A., Deveau, A., Melton, S. J., Kohler, A., Morrell-Falvey, J. L., Brun, A., and Veneault-Fourrey, C. (2014a). Effector MiSSP7 of the mutualistic fungus Laccaria bicolor stabilizes the Populus JAZ6 protein and represses jasmonic acid (JA) responsive genes. Proceedings of the National Academy of Sciences 111:8299–8304.

Plett, J. M., Khachane, A., Ouassou, M., Sundberg, B., Kohler, A., and Martin, F. (2014b). Ethylene and jasmonic acid act as negative modulators during mutualistic symbiosis between L accaria bicolor and P opulus roots. New Phytologist 202:270–286.

Plett, J. M., Yin, H., Mewalal, R., Hu, R., Li, T., Ranjan, P., Jawdy, S., De Paoli, H. C., Butler, G., and Burch-Smith, T. M. (2017). Populus trichocarpa encodes small, effector-like secreted proteins that are highly induced during mutualistic symbiosis. Scientific Reports 7:1–13.

Plett, K. L., Singan, V. R., Wang, M., Ng, V., Grigoriev, I. V., Martin, F., Plett, J. M., and Anderson, I. C. (2020a). Inorganic nitrogen availability alters Eucalyptus grandis receptivity to the ectomycorrhizal fungus Pisolithus albus but not symbiotic nitrogen transfer. New Phytologist 226:221–231.

Plett, J. M., Plett, K. L., Wong-Bajracharya, J., de Freitas Pereira, M., Costa, M. D., Kohler, A., Martin, F., and Anderson, I. C. (2020b). Mycorrhizal effector PaMiSSP10b alters polyamine biosynthesis in Eucalyptus root cells and promotes root colonization. New Phytologist 228:712–727.

Plett, J. M., Kohler, A., and Martin, F. (2024a). Masters of Manipulation: How Our Molecular Understanding of Model Symbiotic Fungi and Their Hosts Is Changing the Face of “Mutualism.” Fungal Associations. The Mycota 9.

Plett, J. M., Wojtalewicz, D., Plett, K. L., Collin, S., Kohler, A., Jacob, C., and Martin, F. (2024b). Sesquiterpenes of the ectomycorrhizal fungus Pisolithus microcarpus alter root growth and promote host colonization. Mycorrhiza 34:69–84.

Plomion, C., Aury, J.-M., Amselem, J., Leroy, T., Murat, F., Duplessis, S., Faye, S., Francillonne, N., Labadie, K., and Le Provost, G. (2018). Oak genome reveals facets of long lifespan. Nature plants 4:440–452.

Qian, M., Xu, L., Tang, C., Zhang, H., Gao, H., Cao, P., Yin, H., Wu, L., Wu, J., and Gu, C. (2020). PbrPOE21 inhibits pear pollen tube growth in vitro by altering apical reactive oxygen species content. Planta 252:1–14.

Radhakrishnan, G. V., Keller, J., Rich, M. K., Vernié, T., Mbadinga Mbadinga, D. L., Vigneron, N., Cottret, L., Clemente, H. S., Libourel, C., and Cheema, J. (2020). An ancestral signalling pathway is conserved in intracellular symbioses-forming plant lineages. Nature Plants 6:280–289.

Reid, D. E., Ferguson, B. J., and Gresshoff, P. M. (2011). Inoculation-and nitrate-induced CLE peptides of soybean control NARK-dependent nodule formation. Molecular Plant-Microbe Interactions 24:606–618.

Replogle, A., Wang, J., Bleckmann, A., Hussey, R. S., Baum, T. J., Sawa, S., Davis, E. L., Wang, X., Simon, R., and Mitchum, M. G. (2011). Nematode CLE signaling in Arabidopsis requires CLAVATA2 and CORYNE. The Plant Journal 65:430–440.

Roy, S., and Müller, L. M. (2022). A rulebook for peptide control of legume–microbe endosymbioses. Trends in Plant Science Advance Access published 2022.

Roy, S., Torres-Jerez, I., Zhang, S., Liu, W., Schiessl, K., Jain, D., Boschiero, C., Lee, H.-K., Krom, N., and Zhao, P. X. (2024). The peptide GOLVEN10 alters root development and noduletaxis in Medicago truncatula. The Plant Journal Advance Access published 2024.

Rustgi, S., Boex-Fontvieille, E., Reinbothe, C., von Wettstein, D., and Reinbothe, S. (2018). The complex world of plant protease inhibitors: Insights into a Kunitz-type cysteine protease inhibitor of Arabidopsis thaliana. Communicative & integrative biology 11:e1368599.

Segonzac, C., and Monaghan, J. (2019). Modulation of plant innate immune signaling by small peptides. Current Opinion in Plant Biology 51:22–28.

Silverstein, K. A. T., Moskal, W. A., Wu, H. C., Underwood, B. A., Graham, M. A., Town, C. D., and VandenBosch, K. A. (2007). Small cysteine-rich peptides resembling antimicrobial peptides have been under-predicted in plants: *Under-predicted cysteine-rich peptides in plants*. The Plant Journal 51:262–280.

Steidinger, B. S., Crowther, T. W., Liang, J., Van Nuland, M. E., Werner, G. D., Reich, P. B., Nabuurs, G.-J., de-Miguel, S., Zhou, M., and Picard, N. (2019). Climatic controls of decomposition drive the global biogeography of forest-tree symbioses. Nature 569:404–408.

Sun, Y., Wu, Z., Wang, Y., Yang, J., Wei, G., and Chou, M. (2019). Identification of Phytocyanin Gene Family in Legume Plants and their Involvement in Nodulation of *Medicago truncatula*. Plant and Cell Physiology 60:900–915.

Tabata, R., and Sawa, S. (2014). Maturation processes and structures of small secreted peptides in plants. Frontiers in plant science 5.

Takahashi, F., Suzuki, T., Osakabe, Y., Betsuyaku, S., Kondo, Y., Dohmae, N., Fukuda, H., Yamaguchi-Shinozaki, K., and Shinozaki, K. (2018). A small peptide modulates stomatal control via abscisic acid in long-distance signalling. Nature 556:235.

Takata, N., Yokota, K., Ohki, S., Mori, M., Taniguchi, T., and Kurita, M. (2013). Evolutionary Relationship and Structural Characterization of the EPF/EPFL Gene Family. PLoS ONE 8:e65183.

Tang, H., Krishnakumar, V., Bidwell, S., Rosen, B., Chan, A., Zhou, S., Gentzbittel, L., Childs, K. L., Yandell, M., and Gundlach, H. (2014). An improved genome release (version Mt4. 0) for the model legume Medicago truncatula. BMC genomics 15:1–14.

Tavormina, P., De Coninck, B., Nikonorova, N., De Smet, I., and Cammue, B. P. (2015). The plant peptidome: an expanding repertoire of structural features and biological functions. The Plant Cell 27:2095–2118.

Tedersoo, L., and Brundrett, M. C. (2017). Evolution of ectomycorrhizal symbiosis in plants. Biogeography of mycorrhizal symbiosis Advance Access published 2017.

Tedersoo, L., May, T. W., and Smith, M. E. (2010). Ectomycorrhizal lifestyle in fungi: global diversity, distribution, and evolution of phylogenetic lineages. Mycorrhiza 20:217–263.

Teufel, F., Almagro Armenteros, J. J., Johansen, A. R., Gíslason, M. H., Pihl, S. I., Tsirigos, K. D., Winther, O., Brunak, S., von Heijne, G., and Nielsen, H. (2022). SignalP 6.0 predicts all five types of signal peptides using protein language models. Nature biotechnology 40:1023–1025.

Thumuluri, V., Almagro Armenteros, J. J., Johansen, A. R., Nielsen, H., and Winther, O. (2022). DeepLoc 2.0: multi-label subcellular localization prediction using protein language models. Nucleic acids research 50:W228–W234.

Tost, A. S., Kristensen, A., Olsen, L. I., Axelsen, K. B., and Fuglsang, A. T. (2021). The PSY peptide family—expression, modification and physiological implications. Genes 12:218.

Tuskan, G. A., Difazio, S., Jansson, S., Bohlmann, J., Grigoriev, I., Hellsten, U., Putnam, N., Ralph, S., Rombauts, S., and Salamov, A. (2006). The genome of black cottonwood, Populus trichocarpa (Torr. & Gray). science 313:1596–1604.

van Der Heijden, M. G., Martin, F. M., Selosse, M.-A., and Sanders, I. R. (2015). Mycorrhizal ecology and evolution: the past, the present, and the future. New phytologist 205:1406–1423.

van Wyk, S. G., Du Plessis, M., Cullis, C. A., Kunert, K. J., and Vorster, B. J. (2014). Cysteine protease and cystatin expression and activity during soybean nodule development and senescence. BMC plant biology 14:1–13.

Vayssières, A., Pěnčík, A., Felten, J., Kohler, A., Ljung, K., Martin, F., and Legué, V. (2015). Development of the poplar-Laccaria bicolor ectomycorrhiza modifies root auxin metabolism, signaling, and response. Plant physiology 169:890–902.

Wang, B., and Qiu, Y.-L. (2006). Phylogenetic distribution and evolution of mycorrhizas in land plants. Mycorrhiza 16:299–363.

Wang, J., Replogle, A. M. Y., Hussey, R., Baum, T., Wang, X., Davis, E. L., and Mitchum, M. G. (2011). Identification of potential host plant mimics of CLAVATA3/ESR (CLE)-like peptides from the plant-parasitic nematode Heterodera schachtii. Molecular plant pathology 12:177–186.

Wang, C., Yu, H., Zhang, Z., Yu, L., Xu, X., Hong, Z., and Luo, L. (2015). Phytosulfokine Is Involved in Positive Regulation of *Lotus japonicus* Nodulation. MPMI 28:847–855.

Wang, C., Velandia, K., Kwon, C.-T., Wulf, K. E., Nichols, D. S., Reid, J. B., and Foo, E. (2021a). The role of CLAVATA signalling in the negative regulation of mycorrhizal colonization and nitrogen response of tomato. Journal of Experimental Botany 72:1702–1713.

Wang, X., Chen, W., Yao, J., Li, Y., Yeboah, A., Zhu, S., and Zhang, Y. (2021b). The evolution and expression profiles of EC1 gene family during development in cotton. Genes 12:2001.

Wang, P., Zhou, J., Sun, W., Li, H., Li, D., and Zhuge, Q. (2023). Characteristics and function of the pathogenesis-related protein 1 gene family in poplar. Plant Science 336:111857.

Wei, H., Liu, G., Qin, J., Zhang, Y., Chen, J., Zhang, X., Yu, C., Chen, Y., Lian, B., and Zhong, F. (2023). Genome-wide characterization, chromosome localization, and expression profile analysis of poplar non-specific lipid transfer proteins. International Journal of Biological Macromolecules 231:123226.

Wen, J., Lease, K. A., and Walker, J. C. (2004). DVL, a novel class of small polypeptides: overexpression alters Arabidopsis development. The Plant Journal 37:668–677.

Wong-Bajracharya, J., Singan, V. R., Monti, R., Plett, K. L., Ng, V., Grigoriev, I. V., Martin, F. M., Anderson, I. C., and Plett, J. M. (2022). The ectomycorrhizal fungus Pisolithus microcarpus encodes a microRNA involved in cross-kingdom gene silencing during symbiosis. Proceedings of the National Academy of Sciences 119:e2103527119.

Wu, K., Qu, Y., Rong, H., Han, X., Tian, Y., and Xu, L. (2022). Identification and expression analysis of the Populus trichocarpa GASA-gene family. International Journal of Molecular Sciences 23:1507.

Wulf, K., Sun, J., Wang, C., Ho-Plagaro, T., Kwon, C.-T., Velandia, K., Correa-Lozano, A., Tamayo-Navarrete, M. I., Reid, J. B., and García Garrido, J. M. (2024). The role of CLE peptides in the suppression of mycorrhizal colonization of tomato. Plant and Cell Physiology 65:107–119.

Xie, H., Zhao, W., Li, W., Zhang, Y., Hajný, J., and Han, H. (2022). Small signaling peptides mediate plant adaptions to abiotic environmental stress. Planta 255:72.

Yamaguchi, K., and Kawasaki, T. (2021). Pathogen-and plant-derived peptides trigger plant immunity. Peptides 144:170611.

Yang, X., Tschaplinski, T. J., Hurst, G. B., Jawdy, S., Abraham, P. E., Lankford, P. K., Adams, R. M., Shah, M. B., Hettich, R. L., and Lindquist, E. (2011). Discovery and annotation of small proteins using genomics, proteomics, and computational approaches. Genome research 21:634–641.

Yu, L., Di, Q., Zhang, D., Liu, Y., Li, X., Mysore, K. S., Wen, J., Yan, J., and Luo, L. (2022). A legume-specific novel type of phytosulfokine, PSK-δ, promotes nodulation by enhancing nodule organogenesis. Journal of Experimental Botany 73:2698–2713.

Zhang, Z., Liu, L., Kucukoglu, M., Tian, D., Larkin, R. M., Shi, X., and Zheng, B. (2020). Predicting and clustering plant CLE genes with a new method developed specifically for short amino acid sequences. BMC genomics 21:1–17.

Zhang, L., Yang, Y., Mu, C., Liu, M., Ishida, T., Sawa, S., Zhu, Y., and Pi, L. (2022). Control of root stem cell differentiation and lateral root emergence by CLE16/17 peptides in Arabidopsis. Frontiers in Plant Science 13:869888.

Zhang, H., Wang, Q., Blanco-Touriñán, N., and Hardtke, C. S. (2024). Antagonistic CLE peptide pathways shape root meristem tissue patterning. Nature plants Advance Access published 2024.

Zhang, Z., Han, H., Zhao, J., Liu, Z., Deng, L., Wu, L., Niu, J., Guo, Y., Wang, G., and Gou, X. (2025). Peptide hormones in plants. Molecular Horticulture 5:7.

Zhou, P., Silverstein, K. A., Gao, L., Walton, J. D., Nallu, S., Guhlin, J., and Young, N. D. (2013). Detecting small plant peptides using SPADA (small peptide alignment discovery application). BMC bioinformatics 14:1–16.

Zhu, F., Deng, J., Chen, H., Liu, P., Zheng, L., Ye, Q., Li, R., Brault, M., Wen, J., and Frugier, F. (2020a). A CEP peptide receptor-like kinase regulates auxin biosynthesis and ethylene signaling to coordinate root growth and symbiotic nodulation in Medicago truncatula. Plant Cell 32:2855–2877.

Zhu, Y., Song, D., Zhang, R., Luo, L., Cao, S., Huang, C., Sun, J., Gui, J., and Li, L. (2020b). A xylem-produced peptide PtrCLE20 inhibits vascular cambium activity in Populus. Plant Biotechnology Journal 18:195–206.

